# Transcription activation is enhanced by multivalent interactions independent of phase separation

**DOI:** 10.1101/2021.01.27.428421

**Authors:** Jorge Trojanowski, Lukas Frank, Anne Rademacher, Pranas Grigaitis, Karsten Rippe

## Abstract

Transcription factors (TFs) consist of a DNA binding and an activation domain (AD) that are considered to be independent and exchangeable modules. However, recent studies conclude that also the physico-chemical properties of the AD can control TF assembly at chromatin by driving a phase separation into transcriptional condensates. Here, we dissected transcription activation by comparing different synthetic TFs at a reporter gene array with real-time single-cell fluorescence microscopy readouts. In these experiments, binding site occupancy, residence time and co-activator recruitment in relation to multivalent TF interactions were compared. While phase separation propensity and activation strength of the AD were correlated, the actual formation of liquid-like TF droplets had a neutral or inhibitory effect on transcription activation. Rather, we conclude that multivalent AD mediated interactions increase the transcription activation capacity of a TF by stabilizing chromatin binding and mediating the recruitment of co-activators independent of phase separation.

## Introduction

The induction of gene expression in eukaryotes involves the binding of transcription factors (TFs) and co-activators at the promoter to induce the assembly of the active RNA polymerase II (Pol II) transcription machinery (Andersson and Sandelin, 2020; Haberle and Stark, 2018; Osman and Cramer, 2020). The vast majority of TFs contain a structurally well-defined DNA-binding domain (DBD) and a separate activation domain (AD) (Frankel and Kim, 1991). The ADs typically comprise intrinsically disordered regions (IDRs) with acidic residues that keep aromatic residues exposed to the solvent (Staller et al., 2018). Synthetic TFs have been successfully constructed by combining DBDs and ADs in a modular manner (Brent and Ptashne, 1985; Chavez et al., 2015; Sadowski et al., 1988). Frequently employed ADs with a particularly high transcription activation capacity are VP16 from a herpes simplex virus protein (Sadowski et al., 1988) and VPR (VP64-p65-Rta), a tripartite synthetic construct that consists of VP64 (4 copies of VP16) fused to the p65 and Rta ADs (Chavez et al., 2015). In addition, it is well established that the TF promoter binding site occupancy *θ* is a key factor that regulates the strength of transcription activation (Bintu et al., 2005). The value of *θ* is determined by the free TF concentration [TF] and the ratio of the kinetic on- and off-rates for binding: *θ* = [*TF*]/([*TF*] + *k*_*off*_/*k*_*on*_). Thus, the target sites become fully saturated if TF concentrations are sufficiently high. In addition, not only binding site occupancy but also TF residence time as given by *τ*_*res*_ = 1/*k*_*off*_ could determine the transcriptional activation capacity (Brouwer and Lenstra, 2019; Callegari et al., 2019; Clauss et al., 2017; Gurdon et al., 2020; Loffreda et al., 2017; Shelansky and Boeger, 2020). The value of *τ*_res_ can become rate limiting for a multi-step activation process if a TF binding event with a certain duration is required to drive a subsequent reaction that induces transcription. The TF chromatin interactions that determine both the binding site occupancy and the TF residence time are thought to be mostly determined by the DBD. However, a number of recent studies showed that TF assembly at chromatin is not limited to direct interactions of the DBD with DNA. The intrinsically disorder region (IDRs) found in TFs like SP1, TAF15, OCT4, β-catenin, STAT3 and SMAD3 as well as transcriptional co-activators like MED1/19, GCN4 and BRD4 and the unstructured C-terminal domain (CTD) of Pol II can drive the formation of so-called transcriptional condensates at enhancers and promoters (Hnisz et al., 2017; Sabari et al., 2020; Shrinivas et al., 2019). One mechanism frequently invoked for this process is liquid-liquid phase separation (LLPS). Above a critical concentration, multivalent interactions of proteins and RNAs that frequently involve IDRs drive the formation of phase separated liquid-like droplets that sequester their constituting components from the surrounding nucleoplasm (Banani et al., 2017; Choi et al., 2020; Shin and Brangwynne, 2017; Uversky, 2021). The assembly of TFs into liquid-like protein droplets at their target sites could enhance transcription (Schneider et al., 2021; Wei et al., 2020) by (i) increasing the local TF concentration at the promoter, (ii) mediating the recruitment of co-activators and/or additional Pol II complexes (*27)*, and (iii) making the TF target search process more efficient (Brodsky et al., 2020; Kent et al., 2020). However, the assembly of transcriptionally active or silenced compartments at chromatin could be also governed by alternative mechanisms including classical (cooperative) chromatin binding, formation of well-defined multi-subunit protein complexes and bridging interactions between distant binding sites (Erdel et al., 2020; Erdel and Rippe, 2018; Frank and Rippe, 2020; McSwiggen et al., 2019a; McSwiggen et al., 2019b; Rippe, 2021). In current studies a comparison of TFs in the droplet state to the same TFs bound to chromatin but without droplet formation is missing, which is crucial to demonstrate a functional role of transcriptional condensates for gene activation.

Here, we have studied a panel of constitutive and light-inducible synthetic TF constructs with dead-Cas9 (dCas9), reverse *tet* repressor (rTetR) and *lac* repressor (LacI) as DBDs and different ADs. We evaluated these synthetic TFs with respect to their activator residence times and their activation capacity with real-time single-cell fluorescence microscopy readouts and assessed the contribution of liquid droplet formation. Striking differences in chromatin bound residence time, RNA production, histone H3 acetylation at lysine 27 (H3K27ac) and BRD4 recruitment between different TF constructs were observed. Furthermore, we link the phase separation propensity of the AD to TF DNA binding properties and activation capacity. Based on our results we conclude that the ability of a TF to engage in multivalent interactions enhances its activation strength. However, we find no evidence that the formation of liquid droplets *per se* would enhance transcription.

## Results

### TF properties affecting transcription initiation are dissected with modular constructs

We studied a range of TF architectures by creating a toolbox of single- and multi-component transcriptional activation complexes (**Fig. 1A, left**). The three DBDs employed were reverse *tet* repressor (rTetR, DNA binding in the presence of doxycycline), *lac* repressor (LacI) and dead-Cas9 (dCas9) with single guide RNAs (sgRNA) binding to *lac*O or *tet*O operator sites. The ADs comprised VP16, p65, Rta, STAT2 and VPR and were associated with these DBD modules via four different approaches: protein fusion constructs (“fusion”), binding of PP7 coat protein (PCP) AD fusions to PP7 RNA loops (Zalatan et al., 2015) engineered into the sgRNA (“loop”), light-induced heterodimer formation between PHR-AD and DBD-CIBN fusion proteins referred to as BLInCR for blue light induced chromatin recruitment (Kennedy et al., 2010; Rademacher et al., 2017) as well as complexes formed by a PP7-sgRNA, tdPCP-CIBN and PHR-AD constructs (“BLInCR-loop”). The CIBN-dCas9-CIBN localizer of the BLInCR-dCas9 system has previously also been used in the LACE system (Polstein and Gersbach, 2015). Our modular toolbox comprised 12 dCas9-based and 14 bacterial repressor-based TF construct combinations. It allowed us to vary DNA binding, self-interaction and activation domain properties for studying how these features affect transcription or other readouts (**Fig. 1A, Fig. S1, Table S1, S2**). As a model system for transcription activation, we used the human U2OS 2-6-3 reporter cell line (Janicki et al., 2004), which carries multiple copies of a reporter gene integrated in tandem at a single locus. Each reporter gene unit contains *lacO* and *tetO* repeats followed by a CMV core promoter and a reporter gene including 24 copies of the MS2 sequence (**Fig. 1A, right**). This cell line model enables time-resolved measurements to follow TF binding and co-factor recruitment in single living cells. In addition, RNA production can be visualized by binding of fluorescently tagged MS2 coat protein (tdMCP) to the MS2 RNA (**Fig. 1A**) (Pankert et al., 2017).

**Fig. 1.**
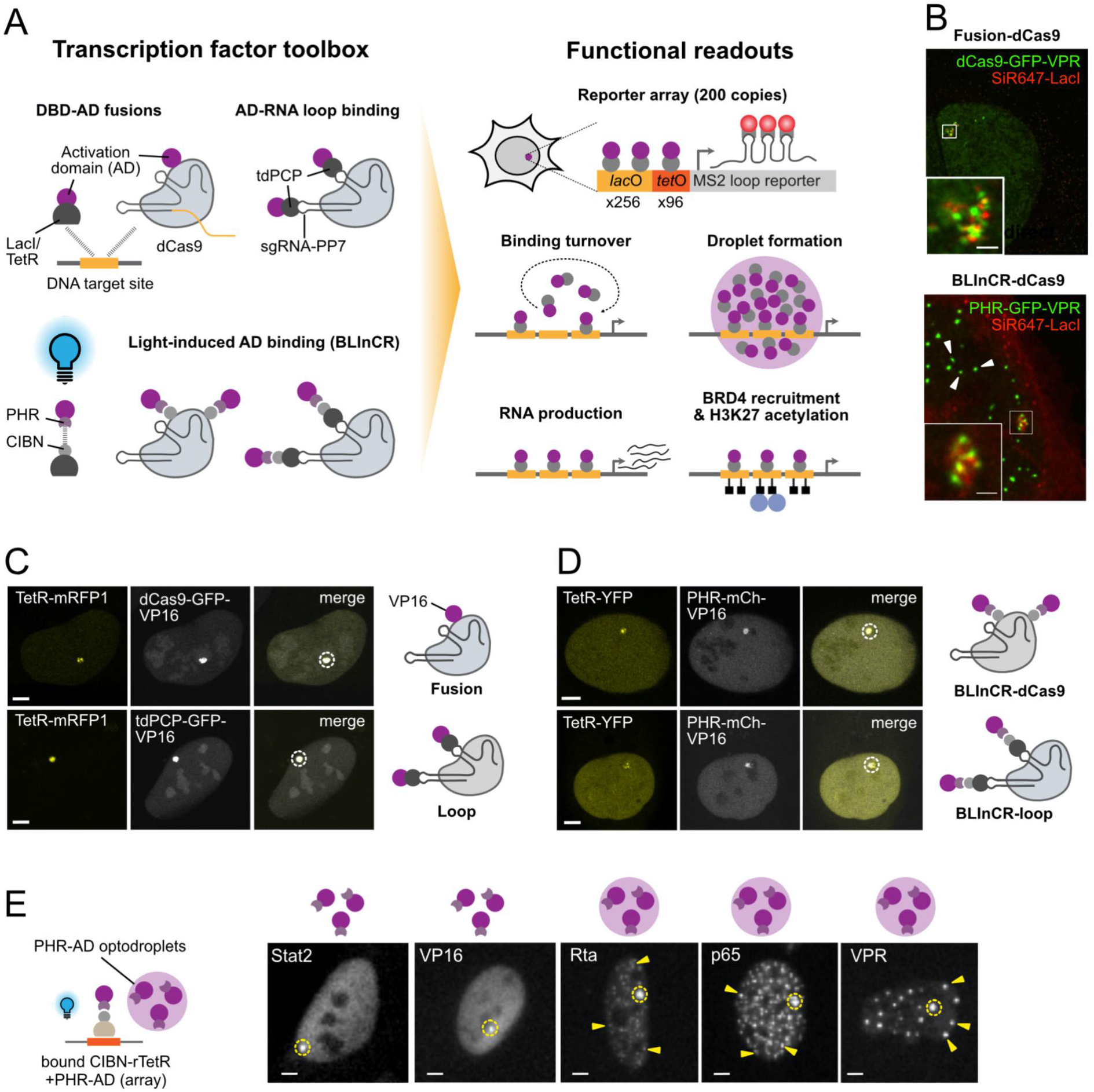
A toolbox to dissect transcription activation. (**A**) (r)TetR, LacI and dCas9 as DBDs were combined with different ADs as direct fusion constructs (“fusion”), via binding of PP7 coat protein to PP7 RNA loops in the single guide RNA (“loop”) or fused to the PHR domain for light induced interactions with CIBN (“BLInCR”). Activity of these TF architectures was analyzed for the different functional readouts depicted. (**B**) SRRF images of the reporter cluster labeled with SNAPtag-LacI/SiR647 and decondensed by dCas9-GFP-VPR fusion (top) or BLInCR-dCas9 VPR (bottom). Scale bar, 1 µm. (**C**) Confocal microscopy images of transfected U2OS 2-6-3 cells showing targeting of the dCas9 complexes to the reporter array marked with a dashed circle. The *tet*O sites of the reporter array were labeled with co-transfected TetR-mRFP1. Scale bars, 5 µm. (**D**) Confocal microscopy images of PHR-mCherry-VP16 enriched at the *lac*O repeats using the BLInCR-dCas9 and BLInCR-loop complexes, respectively. TetR-YFP was used as an array marker. Light-dependency of BLInCR-dCas9/BLInCR-loop complex assembly was confirmed in a separate experiment and compared to CIBN-LacI (**Fig. S1A, S1B**). Scale bars, 5 µm. (**E**) Upon illumination, PHR-GFP-AD binds to CIBN-rTetR upstream of the promoter and can form ectopic optodroplets depending on the AD. Some optodroplets are marked with arrowheads. The reporter array is marked with a dashed circle. Scale bars, 5 µm.

All four types of TF constructs were robustly enriched at the reporter array when using dCas9 with the fluorescently tagged VP16 AD and a sgRNA targeting the *lac*O sites (**Fig. 1C, 1D**). Only a few reporter transcripts were present in untransfected cells and their number increased to thousands of RNA molecules in cells as shown by RNA-FISH for the strong activator VPR (**Fig. S1 D-E**). Separated *lac*O and *tet*O site clusters were visible in super-resolution radial fluctuations (SRRF) microscopy (Gustafsson et al., 2016) images at the activated reporter (**Fig. 1B, top; Fig. S1C**). The use of light-controllable PHR/CIBN modules in our BLInCR type TFs with dCas9 or LacI/rTetR as DBDs enabled fast and reversible binding to dissect transcription initiation at high temporal resolution (**Fig. S1A, B**) (Rademacher et al., 2017). Moreover, PHR constructs can self-associate to form so-called optodroplets under appropriate conditions (Shin et al., 2017) as visualized for PHR-VPR recruited via BLInCR-dCas9 (**Fig. 1B bottom**). These optodroplets were detectable as nuclear punctae outside the reporter array. The granular signal at the reporter indicated that the reporter cluster itself was not immersed in a single homogenous compartment under these conditions. PHR optodroplet formation can be exploited to evaluate the phase separation propensity of TF ADs in living cells (Erdel et al., 2020; Shin et al., 2017). BLInCR-mediated recruitment of different AD fusions to the *tetO* sites of the reporter revealed that optodroplets formed to a very different extent (**Fig. 1E**). Rta, p65 and VPR readily assembled into optodroplets whereas they were rare or absent for VP16 and STAT2. We quantified optodroplet formation propensity of the constructs depicted in **Fig. 1E** by manually determining the fraction of cells with visible optodroplets, which ranged from <1% (STAT2) and 29% (VP16) to 41% (Rta), 72% (p65) and 86% (VPR) (**Table S3**). We also determined the area of optodroplets relative to the nuclear area as an additional measure of droplet abundance in dependence of the nuclear concentration (**Fig. S2A**). From this relationship, we deduced critical concentrations for droplet formation as the nuclear intensities at which the relative droplet area crossed an empirically defined threshold. These critical concentrations (in a. u.) ranged from 0.19 for VPR with a high self-interaction propensity to 0.54 (VP16) and >1.5 (STAT2) (**Table S3**). Thus, the propensity to form droplets of the ADs used in our toolbox covered a broad range, enabling us to link this feature to other readouts of the reporter system.

### Activation strength correlates with phase separation propensity

Phase separation has been used to explain the assembly of TFs into transcriptionally active compartments and was suggested to play a role in amplifying transcription (Hnisz et al., 2017; Sabari et al., 2018; Schneider et al., 2021; Shrinivas et al., 2019; Wei et al., 2020). To challenge this model, we used BLInCR TF constructs to directly correlate the phase separation propensity of our ADs with their potential to induce transcription of the reporter array by monitoring nascent RNA production with tdMCP-tdTomato (**Fig. 2A**). The ectopic AD assemblies displayed properties indicative of liquid droplets like fusion, higher mobility of droplets compared to the reporter array (**Fig. S2B**) and predominantly fast exchange with the nucleoplasm as determined by fluorescence recovery after photobleaching (FRAP) (**Fig. S2C**). To link phase separation propensity to transcriptional activation the ADs were recruited to the reporter and single cell RNA production trajectories were recorded (**Fig. 2A**). All ADs were able to elicit a transcriptional response and the activator strength was positively correlated with their propensity to form optodroplets (**Fig. 2B-D, Table S3, S4**): (i) In the first 90 minutes of transcription activation, p65, Rta and VPR displayed higher maximum transcription levels of 1.7-2.9 a. u. compared to 1.3-1.6 a. u. for VP16 and STAT2 (*p* < 0.001, two-way ANOVA) (**Fig. 2B, S2D**). (ii) The fraction of responding cells after 90 minutes was larger with 67-92% (p65, Rta, VPR) vs. 42-67% (VP16, STAT2) (**Fig. 2C**). (iii) The time to reach half-maximal activation was shorter with 26-28 min (p65, Rta, VPR) compared to 38-42 min (VP16, STAT2) (**Table S4, Fig. S2E**) and increased with the critical concentration for droplet formation (**Fig. 2D**). Thus, the strong transcriptional activators VPR, p65 and Rta showed a high phase separation propensity while VP16 and STAT2 were weaker activators with a low tendency to form liquid droplets.

**Fig. 2.**
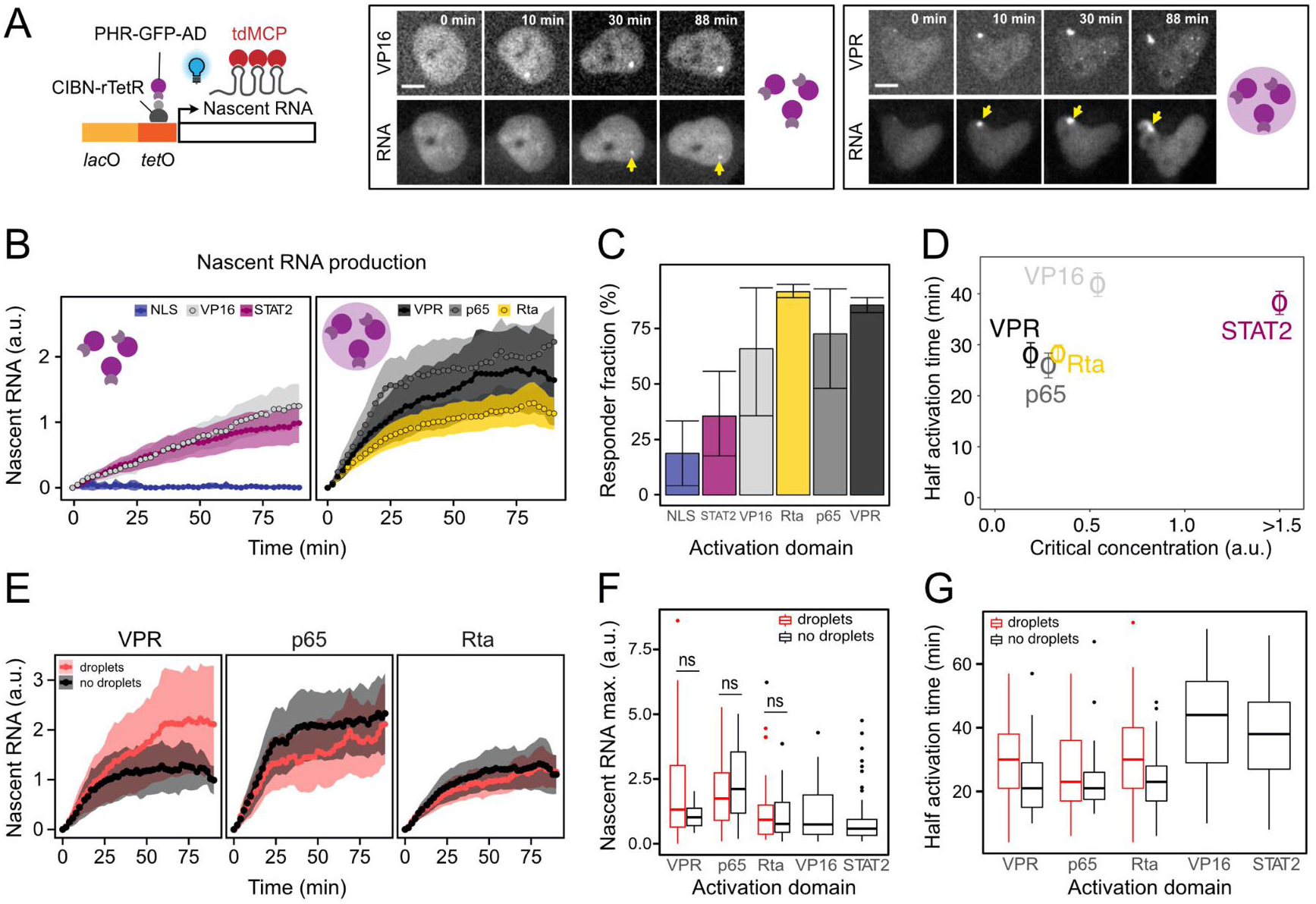
AD phase separation propensity and activation capacity. Single cell time courses of nascent RNA production were acquired to investigate the relation between droplet formation and transcription activation. PHR-GFP-AD to CIBN-rTetR (BLInCR-rTetR) binding was triggered by blue light illumination and optodroplet formation was monitored in the PHR-GFP-AD channel. (**A**) Experimental approach and representative image series for ADs with high (VPR) and low (VP16) droplet forming propensity. Scale bars, 10 µm. (**B**) Average nascent RNA kinetics of responding cells for the different ADs with mean and 95% CI and *n* = 31-71 cells per construct. PHR-GFP-NLS served as negative control. (**C**) Fraction of cells with visible enrichment of tdMCP at the reporter (**Table S4**). Bars represent minimum and maximum values from 2 or 3 replicate experiments per construct. (**D**) Plot of activation speed against phase separation propensity. The time to half maximal activation was determined from the single cell trajectories. Critical concentrations for optodroplet formation were determined from microscopy images (**Fig. 1E, S2A**). Error bars represent s. e. m. values. (**E**) Average activation time courses of responding cells visually classified as droplet containing or not with 95% CI (*n* = 13-18 cells per condition). (**F**) RNA production plateau values for different ADs calculated from the last five time points. For VPR, p65 and Rta cells with (red) or without (black) optodroplet formation outside the reporter array were compared. Per condition *n* = 13-55 cells were analyzed while excluding non-responding cells. n. s., not significant (*p* > 0.05, unpaired two-sided Welch’s *t*-test). (**G**) Time to half maximal activation of samples described in panel F. The presence of droplets had a neutral or slightly inhibitory effect (*p* = 0.09, two-way ANOVA).

### Phase-separated TF compartments are not required for efficient transcription

The nascent RNA time course data were split into cells with and without visible optodroplets to elucidate the relation between droplet formation and transcriptional activation. The presence of PHR-GFP-AD optodroplets in a given cell indicated that the conditions in this cell were above the critical threshold for phase separation, implying that also the reporter gene cluster was in a phase-separated environment. The comparison of transcription activation kinetics between the two groups showed no enhanced activation rate for time courses that displayed optodroplet formation (**Fig. 2E**). There was no significant difference in the maximum value of RNA production for cells with and without droplet formation (*p* > 0.05 in pairwise *t*-test and in two-way ANOVA) (**Fig. 2F, Table S4**). If droplets were present, the time to half activation was unchanged for p65 (26 ± 8 min) and even moderately increased for VPR (from 25 ± 6 min to 30 ± 9 min) and Rta (from 25 ± 4 min to 31 ± 5 min) (0.05 ≤ *p* ≤ 0.10, two-way ANOVA accounting for AD type and presence/absence of droplets) (**Fig. 2G, Fig. S2F**). We conclude that droplet formation does not enhance transcription activation but rather displayed a trend to a moderate inhibition.

Next, we tested if enhancing the intrinsically low droplet formation propensity of VP16 (**Fig. 1E, S2A**) affects transcription activation by applying three different approaches for BLinCR rTetR-mediated recruitment of VP16 (**Fig. 3A**): (i) Co-transfection with CIBN-LacI, which can act as a bridging factor between PHR-AD molecules via interactions between LacI dimers (Lewis et al., 1996). (ii) Increasing the droplet formation propensity by binding of a second PHR domain to the AD via a GFP binding protein (GBP) construct fused to PHR, which will generate VP16 complexes with two PHR domains via the high affinity GFP-GBP interaction. (iii) Fusion of VP16 to the N-terminal IDR of the FUS (fused in sarcoma) protein (FUSN), which has a high propensity to form liquid droplets *in vitro* and *in vivo* (Patel et al., 2015). In the first approach, the addition of CIBN-LacI enhanced droplet formation and increased the VP16 concentration around the promoter (**Fig. 3B, top row, Fig. S3A, B**). This was in line with our observation that recruiting PHR-GFP-VP16 to CIBN-LacI instead of CIBN-TetR already lowered the critical concentration for droplet formation (**Fig. S2A, Table S3**). Surprisingly, nascent RNAs levels at the reporter were largely reduced by addition of CIBN-LacI (**Fig. 3C-E, Fig. S3C-D**). The fraction of responding cells decreased from 70 to 24% and a more than 2-fold reduction of bulk RNA levels was measured by quantitative real-time PCR (qPCR). Co-transfection of the non-bridging GFP-LacI control instead of CIBN-LacI revealed some repression by GFP-LacI binding alone, but this effect was much smaller than the repression observed by CIBN-LacI induced optodroplets. In a second set of experiments, we increased the valency of PHR-GFP-VP16 by co-transfection with PHR-GBP that binds to it. This approach was highly effective in inducing VP16 droplet formation (**Fig. 3F, Fig. S3E**). However, transcription was almost complete repressed as apparent from the maximum nascent RNA level and the fraction of responding cells (**Fig. 3G-H, Fig. S3F**). Measurements of bulk reporter RNA levels by qPCR confirmed this conclusion (**Fig. 3I)**. The third approach induced VP16 droplets by fusion to FUSN (**Fig. S3A, E)**. This PHR-GFP-FUSN-VP16 construct displayed a largely increased activation strength, which was similar to that of VPR in the measurements of transcription kinetics (**Fig. 3G-H, Fig. S3F**). In addition, bulk RNA levels were 10-fold higher than those measured for VP16 without FUSN (**Fig. 3I**). FUSN alone failed to activate the reporter. Next, we split up the FUSN-VP16 nascent RNA time course data into cells with and without visible droplets. Notably, we found that activation capacity of FUSN-VP16 was indistinguishable between the two groups (**Fig. S3H**). This is in agreement with the results showing that actual droplet formation had no effect on the activation capacity of VPR, p65 and Rta (**Fig. 2E**). We conclude that multivalent interactions increase activation capacity but that the additional formation of droplets has no effect on transcriptional activation. Moreover, certain types of droplets like those containing bivalent bridging factors can efficiently inhibit transcription activation even though they increase the local TF concentration at the promoter.

**Fig. 3.**
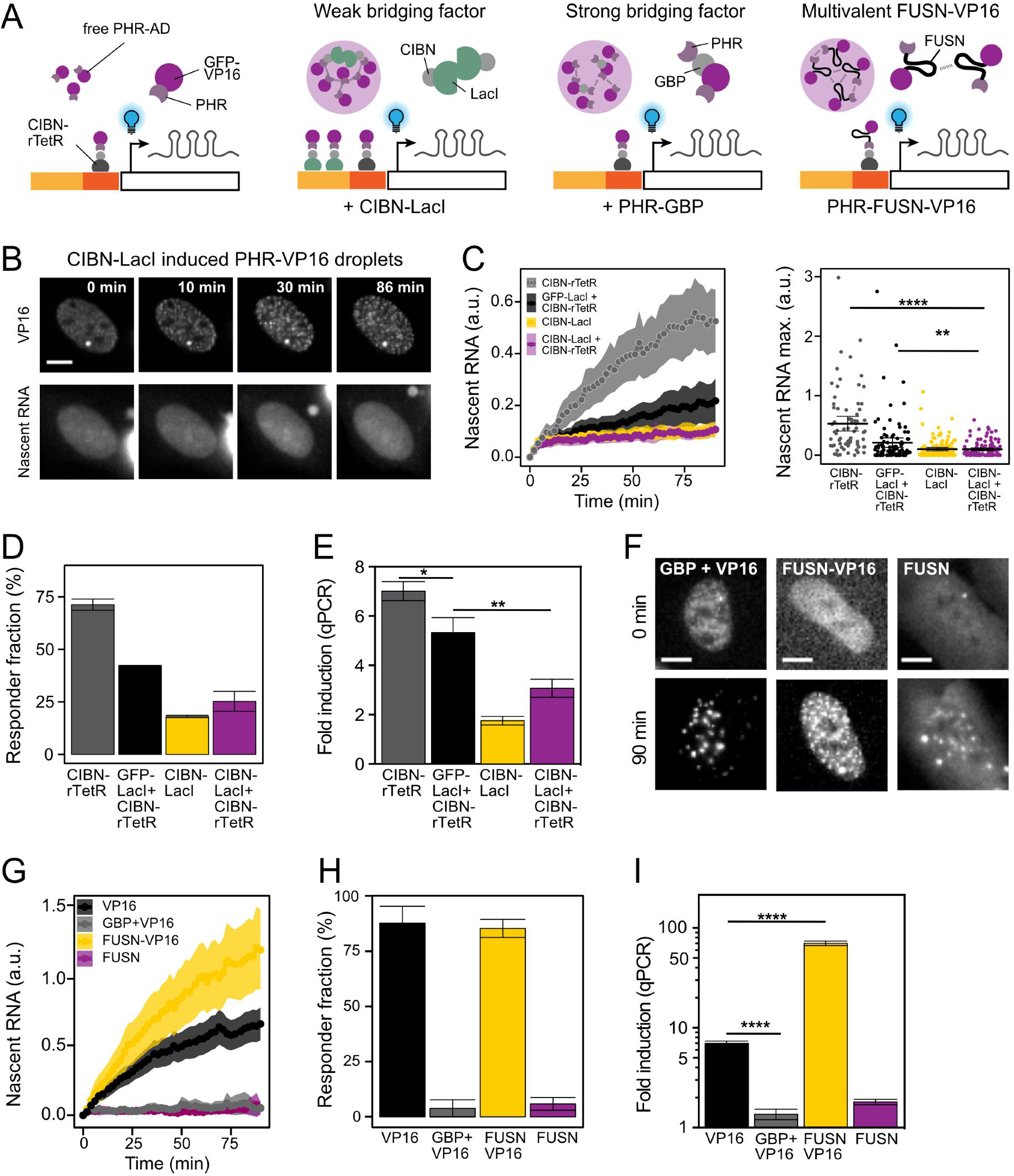
Effects of induced droplet formation of VP16 on transcription activation. (**A**) Three independent approaches to increase the droplet formation propensity of VP16 were employed. The effect on transcription was studied for both single cell nascent RNA and bulk RNA production. (**B**) Microscopy images of a nascent RNA production time course showing PHR-GFP-VP16 optodroplet induction by CIBN-LacI co-transfection with CIBN-rTetR in a non-responding cell. (**C**) Left: Averaged nascent RNA time courses and 95% CI for activation in presence of CIBN-LacI induced optodroplets including responding and non-responding cells (*n* = 74-126 cells per condition). Right: The maximum values reached at the time course plateau. The activation potential is reduced by CIBN-LacI with CIBN-rTetR for VP16 as compared to the unperturbed state with only CIBN-rTetR and the GFP-LacI co-transfection control. Dots, single cell RNA time course maxima; bar, mean; error bars, 95% CI; n. s., not significant, *p* > 0.05; *, *p* < 0.05; **, *p* < 0.01; ****, *p* < 0.0001. The *p* values were computed with a two-sided Welch’s *t*-test. (**D**) Fraction of cells with visible enrichment of tdMCP at the promoter spot. Error bars represent minimum and maximum of 2 experiments per condition. (**E**) qPCR of bulk reporter RNA levels 90 min after induction. Data are represented as mean and s. d. of the fold-change induction compared to mock transfected samples and normalized to beta actin mRNA (*n* = 3). *, *p* < 0.05; **, *p* < 0.01 from an unpaired two-sided Student’s *t*-test. (**F**) Image series of cells at the beginning and end of time courses showing optodroplet formation for PHR-GFP-VP16 plus PHR-GBP, PHR-GFP-FUS-VP16 and PHR-GFP-FUS (control). Scale bar: 10 µm. (**G**) Average nascent RNA time courses and 95% CI for activation by GBP and FUSN induced optodroplets including responding and non-responding cells. PHR-GFP-FUSN was used as control. *n* = 24-154 cells per condition. (**H**) Fraction of cells with visible enrichment of the tdMCP labeled RNA at the array for GBP/FUSN experiments. Error bars represent minimum and maximum of 2 experiments per condition. (**I**) Bulk reporter RNA levels measured by qPCR at 90 min after induction for GBP/FUSN experiments. Data represent mean and s. d. of fold-change induction compared to mock transfected samples and normalized to beta actin mRNA (*n* = 3). *p* < 0.0001 (****) calculated from an unpaired two-sided Student’s *t*-test.

### TF residence times are determined by both the DBD and the AD

Our experiments with the BLInCR TFs suggest that multivalent interactions of ADs outside the phase separation concentration regime are crucial for efficient transcription activation. One possible mechanism could be to stabilize chromatin binding and to increase TF residence time. Accordingly, we conducted a FRAP analysis of different TF constructs containing either an AD with high (VPR) or low (VP16) multivalent interaction potential (**Fig. 4A-C, S4A-D**). The recovery curves obtained by bleaching the TFs bound at the reporter gene array were fitted by a reaction-diffusion model for clustered binding sites as described in the **Supplementary Methods** (Sprague et al., 2006). This analysis yielded the apparent diffusion coefficient, the dissociation rate *k*_off_ and the immobile fraction of stably bound molecules during the observation period of 240 s (**Fig. S4E, Table S5**). The computed residence times in the chromatin bound state ranged from *τ*_res_ = 12 ± 6 s for tdPCP-GFP binding to PP7-loops of the dCas9-sgRNA to a large immobile fraction with *τ*_res_ >240 s for the dCas9-GFP-VP16 fusion construct (**Table S5, Fig. 4C**). Interestingly, the residence times for VPR as compared to VP16 were consistently >14 s higher and the immobile fraction was increased by 2-7% in the highly dynamic loop and BLInCR complexes (**Fig. 4C**). This observation points to VPR self-interactions that stabilize the binding of the tdPCP-GFP-VPR and PHR-GFP-VPR constructs. For the already very stably chromatin bound dCas9 fusion constructs the VPR-VPR interactions manifested themselves as an additional faster exchanging contribution to the recovery that is clearly visible for dCas9-GFP-VPR but absent in dCas9-GFP-VP16 (**Fig. 4C, Table S5**). To confirm this additional indirect protein binding at the array, we measured the intensity of VPR and VP16 assemblies recruited via the loop configuration at the reporter array (**Fig. 4D**). We found a 1.9-fold higher signal for tdPCP-GFP-VPR as compared to tdPCP-GFP-VP16 (*p* = 0.0006, Welch two-sample *t*-test), confirming the additional recruitment seen in FRAP (**Fig. 4C, upper row third panel**). We conclude that the TF residence time was not only dependent on the DBD but significantly influenced by the AD properties. VPR promoted the binding of additional TF molecules to those that were directly bound to DNA and stabilized the directly bound proteins in the case of weaker interactions. This enrichment of TFs via protein-protein interactions correlated with the AD’s propensity to engage in multivalent interactions.

**Fig. 4.**
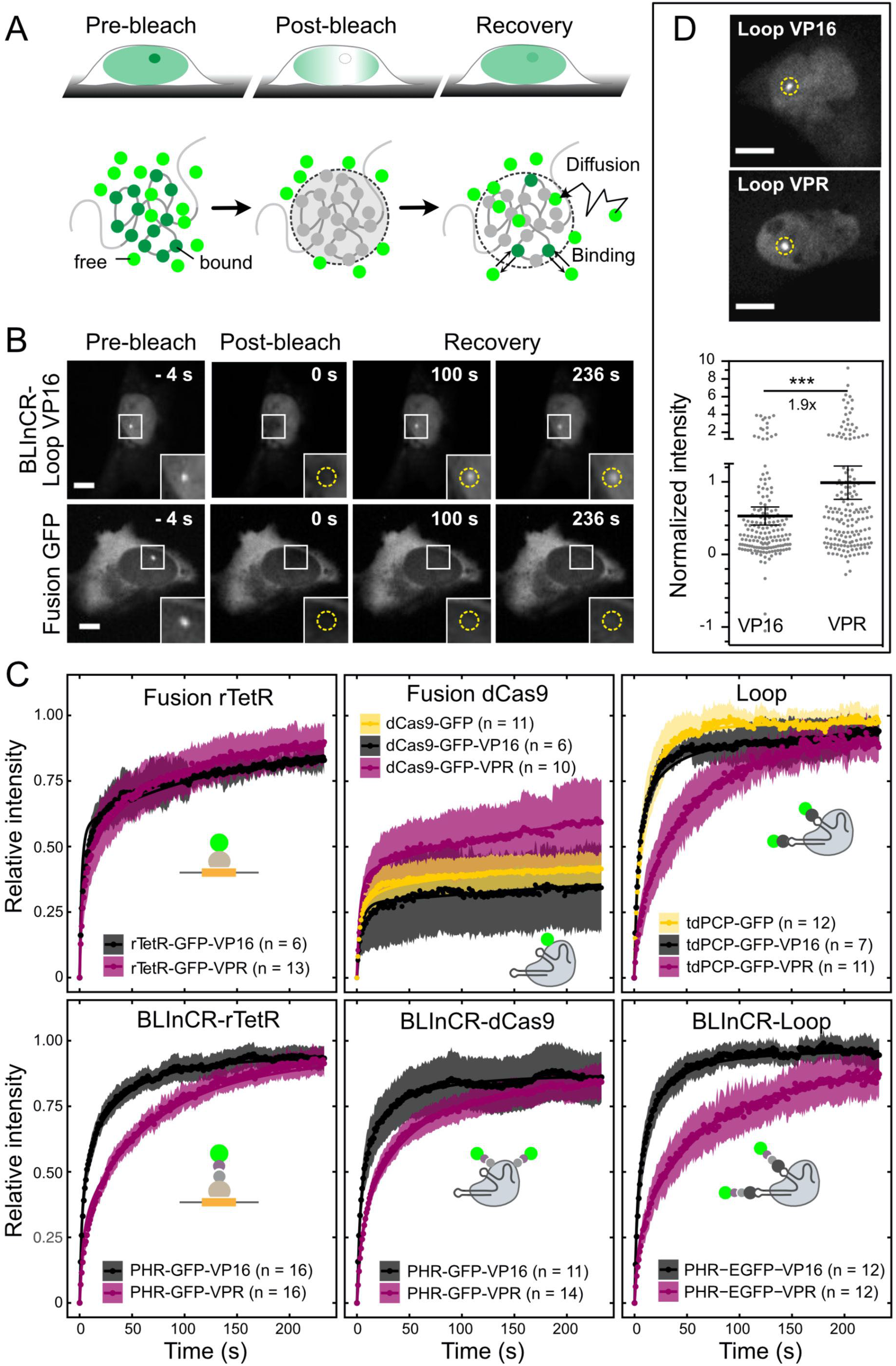
FRAP measurements of different TF complexes. (**A**) Experimental setup for FRAP reaction-diffusion analysis of TF binding. After bleaching fluorescence recovers by both diffusion of unbleached molecules and exchange of bound molecules at the promoter binding sites in the U2OS 2-6-3 reporter gene array. (**B**) Exemplary FRAP image series for the BLInCR-loop VP16 complex with fast exchange (top) and for dCas9-GFP fusion complex displaying no exchange of DNA bound molecules during the observation period (bottom). Scale bar: 10 µm. The dashed circle in the inset marks the reporter array. (**C**) Averaged FRAP curves and 95% CI. Constructs were recruited to *lacO* except for the *tetO*-dependent constructs. The solid line represents the fit of the data to a reaction-diffusion model for clustered binding sites for GFP-tagged complexes of VP16 and VPR with the indicated DNA binding modules. (**D**) Enrichment of tdPCP-GFP-VP16 and tdPCP-GFP-VPR in the loop complex at the reporter array. GFP signal was background subtracted and normalized to tagBFP-LacI as a marker of the binding site cluster. The 1.9-fold higher signal for VPR indicates an increased amount of indirectly bound molecules. Solid bar, mean; error bars, 95% CI; *n* = 164-166 cells per condition; *p* < 0.001 (***) calculated from unpaired two-sided Welch’s *t*-test. Scale bar: 10 µm.

### Transcription activation can occur independent of BRD4 and H3K27ac

TFs initiate transcription via different mechanisms that include the assembly of the transcription machinery, catalyzing its transition into an active state as well as chromatin state changes that promote transcription (Lee and Young, 2000). In order to unravel how these mechanistic aspects are linked to the multivalent interaction and/or the residence time of a TF, we established an end-point assay to correlate activity of TF constructs studied with marks of active transcription (**Fig. 5**). We first measured the activity of the TF constructs by qPCR of bulk reporter RNA levels 24 hours after induction (**Fig. 5A, Table S6**). In these experiments, the BLInCR-dCas9 and BLInCR-loop constructs failed to activate the core CMV promoter of the gene array. Transcriptional activation by a construct similar to BLInCR-dCas9 has been reported previously for the endogenous human *IL1RN* or *HBG1/2* promoters and to a lesser extent for *ASCL1* (Polstein and Gersbach, 2015). This apparent discrepancy was further studied as described in the **Supplemental Information** (**Fig. S5A-C**) and is likely to reflect differences of the promoters studied. All other TF constructs were able to activate transcription. VPR was a stronger activator than VP16 in most cases, as shown also by the steady-state nascent RNA profiles determined from the tdMCP signal after 24 hours (**Fig. 5D**). Additionally, BRD4 and H3K27ac were measured by fluorescence microscopy (**Fig. 5B-C**) to yield their normalized radial profiles across the gene array (**Fig. 5D**). This analysis revealed that VPR displayed much stronger BRD4 and H3K27ac enrichment than VP16 except for the loop construct. As an internal control, we also recruited the histone acetyltransferase p300 core domain fused to dCas9. The binding in p300 led to H3K27ac and BRD4 accumulation but not to RNA production. Notably, the transcriptionally inactive BLInCR-dCas9 and BLInCR-loop complexes were capable to induce enrichment of both BRD4 and H3K27ac (**Fig. 5D, Supplemental Information**). We conclude that transcription activation can occur in parallel to co-activator recruitment and histone acetylation and distinguish three different TF types: (i) Strong activators like dCas9-VPR and rTetR-VPR fusions that induce both transcription and strong enrichment of BRD4 and H3K27ac. (ii) Activators represented by the rTetR-VP16 fusion that displayed moderate but robust activation at very low levels of BRD4 and H3K27ac. (iii) The dCas9-based BLInCR constructs that efficiently recruited BRD4 and induced H3K27 acetylation but failed to activate transcription.

**Fig. 5.**
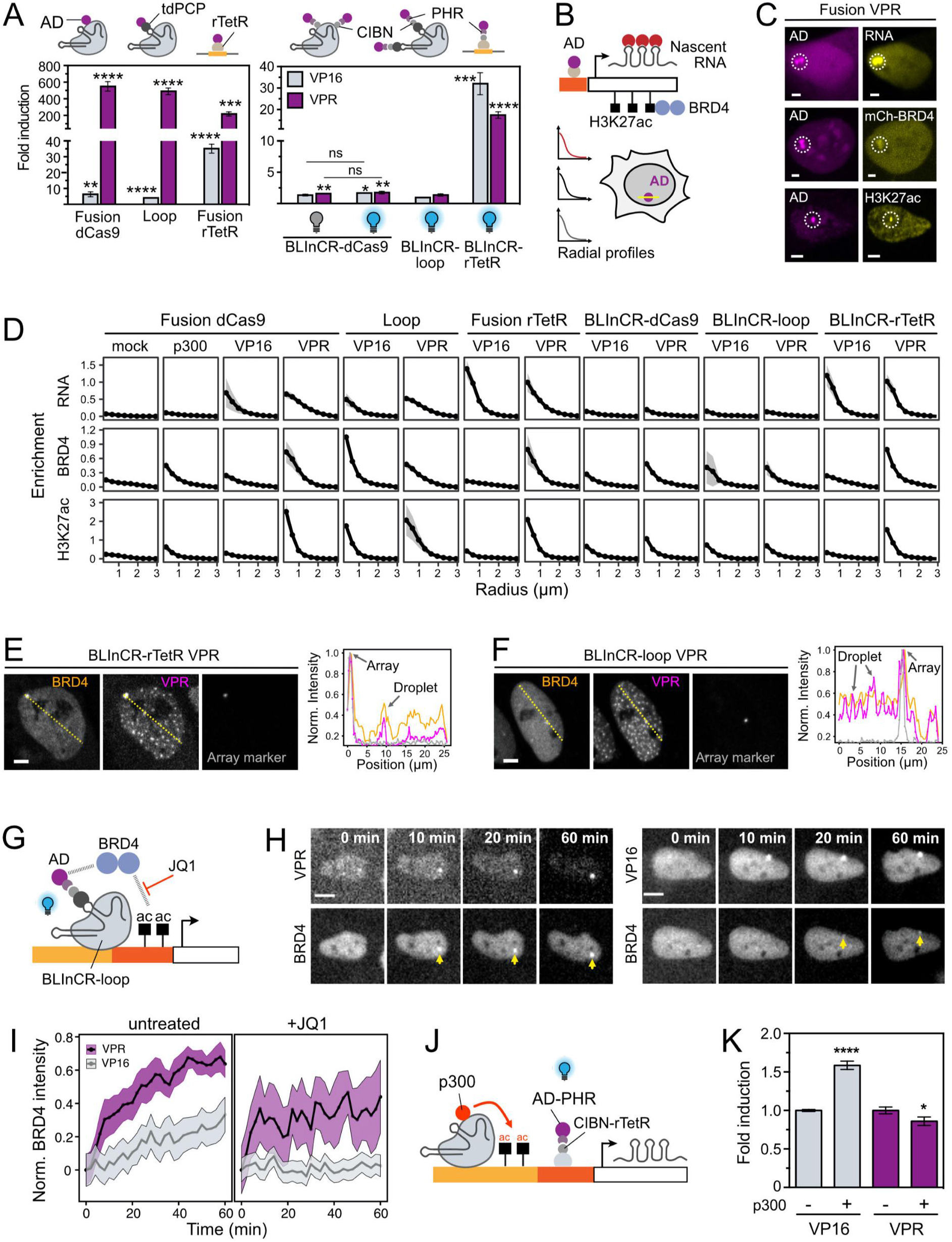
Transcription activation features in dependence of TF architecture and AD type. (**A**) Quantification of reporter RNA by qPCR for the different TF constructs recruited to the *tetO* sites at 24 hours after transfection and induction by addition of doxycycline and/or constant blue light illumination. Data represent mean and s. d. of fold-change upon induction compared to mock transfected samples and normalized to beta actin mRNA (*n* = 3). Indicated *p*-values of > 0.05 (n. s.), < 0.05 (*), < 0.01 (**), < 0.001 (***), < 0.0001 (****) are from a two-sided unpaired Student’s *t*-test against the mock condition. (**B**) Enrichment analysis of activation marks and nascent RNA in single cells 24 h after induction. Activators were recruited to the *tet*O sites and radial enrichment profiles were measured by confocal microscopy of living (nascent RNA, mCherry-BRD4) or fixed (H3K27 acetylation immunostaining) cells. (**C**) Representative microscopy images of cells with *tet*O-targeted dCas9-GFP-VPR complex and the three readouts: nascent RNA visualized with co-transfected tdMCP-tdTomato, co-transfected mCherry-BRD4 and immunostaining against H3K27ac. The dashed line circle marks the reporter array. Scale bar: 5 µm. (**D**) Radial enrichment profiles for RNA, BRD4 and H3K27ac for all TF architectures with either VP16 or VPR. Profiles depict the average of *n* = 16-520 cells per condition. A dCas9-GFP (mock) and dCas9-GFP-p300core fusion were included as additional reference conditions. (**E**) Cells showing partial co-localization of overexpressed mCherry-BRD4 with PHR-GFP-VPR optodroplets upon recruitment via BLInCR-rTetR. Co-transfected tagBFP-LacI marks the reporter array. Normalized intensity profiles showing the enrichment of mCherry-BRD4 (orange) in some of the VPR optodroplets (pink) and at the array spot marker (grey). Intensities were normalized to the maximum value within each channel profile. Profile positions are indicated by dashed lines. Scale bar, 5 µm. **(F)** Same as panel E but for the BLInCR-loop construct. (**G**) Experimental setup for BRD4 co-recruitment by different activators. Light-induced binding of the activator to the BLInCR-loop complex on both *tet*O and *lac*O sites with or without the bromodomain inhibitor JQ1. The complex does not induce transcription and thus indirect effects of the active transcription machinery on BRD4 binding/acetylation are absent. (**H**) Representative live cell image time series of mCherry-BRD4 enrichment at the reporter (arrows) for VPR (left) or VP16 (right) in the absence of JQ1. Scale bar, 10 µm. (**I**) Time traces of BRD4 signal accumulation at the reporter after light induced VPR/VP16 binding without JQ1 (left) or with JQ1 treatment (right) starting three hours before light-induction. Mean values of normalized intensity and 95% CI are shown with *n* = 10-85 cells per condition. (**J**) Experimental setup to test VP16 and VPR activity in the presence of pre-existing histone acetylation. A dCas9-GFP-p300 fusion was constitutively recruited to the *lac*O sites to induce acetylation prior to transcription activation by VPR or VP16 BLInCR-rTetR for 90 minutes. (**K**) qPCR measurements of reporter RNA levels for the experiment outlined in (**J**). Fold induction of reporter RNA for the dCas9-GFP-p300 (+) relative to dCas9-GFP (-) control condition is represented as mean and s. d. of 3 replicates. Data was normalized to beta actin mRNA levels and to the reporter RNA level of the dCas9-GFP condition (-). A two-sided unpaired Student’s *t*-test between p300 (+) and control (-) was used to calculate *p*-values of < 0.05 (*) and < 0.0001 (****).

### VPR recruits BRD4 directly and is less dependent on pre-existing histone acetylation

We observed that both activating and non-activating VPR BLInCR complexes sometimes enriched over-expressed mCherry-BRD4 in ectopic optodroplets while this was hardly observed for VP16 (**Fig. 5E, F**). Thus, we hypothesized that transient multivalent interactions between VPR and BRD4 could contribute to the faster (**Fig. 2B**) and often stronger (**Fig. 5A**) transcription activation by VPR compared to VP16. To test this possibility, we used the transcription-incompetent BLInCR-loop complex to monitor transcription-independent BRD4 binding after light-induced recruitment of VP16 or VPR (**Fig. 5G**). BRD4 accumulation was strong and fast for VPR with an initial steep rise of the BRD4 levels over the first 10 minutes followed by a phase of slower BRD4 accumulation (**Fig. 5H-I**). VP16 did not display such biphasic kinetics. Next, we conducted the same experiment after treating the cells with the inhibitor JQ1 that disrupts BRD4 interactions with acetylated histones to evaluate the contribution of BRD4 binding that is independent of acetylation. JQ1 pre-treatment completely abrogated BRD4 accumulation for VP16. For VPR, the initial steep rise remained unaffected, but BRD4 binding was reduced in the second phase (**Fig. 5I, right**). This observation suggests a direct VPR-BRD4 interaction that is followed by subsequent binding of BRD4 to acetylated histones via its bromodomain.

Next, we examined the active BLInCR-rTetR complex to study the role of BRD4 for transcription. With this complex, JQ1 treatment did not reduce nascent RNA production induced by VP16 or VPR (**Fig. S5D, Table S7**). Bulk RNA levels were moderately reduced in the qPCR assay for VPR (1.8-fold reduction) but mostly unaffected for VP16 (1.1-fold reduction) (**Fig. S5E, Table S6**). We conclude that BRD4 accumulation accompanies transcriptional activation but is not essential for reporter gene induction. It may, however, enhance transcription activation by VPR or stabilize the activated state. Next, we assessed the contribution of histone acetylation to the activation capacity of both VP16 and VPR. The dCas9-p300core construct was constitutively recruited to the distal *lac*O sites. It establishes a locally hyper-acetylated state without inducing transcription (**Fig. 5D**). Subsequently, light-induced transcription by VP16 and VPR BLInCR-rTetR was compared (**Fig. 5J**). VP16 displayed an enhanced transcriptional response quantified by qPCR in the presence of p300 at the *lac*O repeats compared to the case without p300 whereas VPR did not (**Fig. 5K, Table S6**). When observing nascent RNA at the gene array by microscopy, a more pronounced increase of the plateau level was observed for VP16 (3.1-fold) than for VPR (2.1-fold) while activation kinetics were similar (**Fig. S5G, Table S7**). We conclude that pre-existing histone acetylation can increase the transcription induction and does so to a higher extent for VP16. Accordingly, the lower transcription induction of VP16 could be related in part to its inability to directly interact with histone acetylases like p300 as well as the BRD4 co-activator.

### Shortened residence times reduces activation capacity

The preceding comparison of different TF complexes revealed links between phase separation propensity, TF turnover and transcriptional activation capacity. The impact of TF turnover on transcription was clearly dominated by the specific TF complex architecture (e.g., loop vs direct dCas9 fusion). To directly dissect how binding affinity and residence time affect transcriptional output, we artificially increased the turnover of DNA-bound dCas9 complexes at the promoter by introducing a C2G mutation into the sgRNA targeting the *tet*O sequence to yield the sgRNA-mut (**Fig. 6A, Fig. S6A, Table S2**). The FRAP analysis showed that the residence time of the dCas9 VPR fusion construct for sgRNA-mut versus sgRNA-wt was strongly decreased from *τ*_res_ = 124 s (95% CI: 75-347 s) to *τ*_res_ = 57 s (95% CI: 34-184 s) (**Fig. 6B, Table S5**). In addition, the immobile fraction was lowered from 36% to 7% (95% CIs: 25-47% and 0-16%). Next, we compared transcription activation with sgRNA-mut versus sgRNA-wt. For sgRNA-mut complexes, occupancy was reduced to 14% (VP16) and 37% (VPR) of the respective sgRNA-wt value (**Fig. 6C, 6D**). The fraction of cells with visible dCas9 recruitment decreased from 90% to 76% (VPR) and from 59% to 17% (VP16) (**Fig. S6B, Table S8**) and bulk reporter RNA levels were strongly reduced for both VP16 and VPR (**Fig. 6E**). For sgRNA-wt, a continuous increase of occupancy with activator expression was observed, which is likely to reflect an increase of indirectly recruited molecules via multivalent interactions. As expected, RNA production increased with occupancy. In order to separate the effect of occupancy and residence time, we binned cells into groups with equal occupancy and compared their nascent RNA production. The average RNA production induced by VPR was consistently lower for the shorter residence times of sgRNA-mut vs. the longer residence times of sgRNA-wt within each TF occupancy group (**Fig. 6F**) (VPR: 2 - 6-fold, VP16: 1.3 - 2-fold; VPR: *p* < 0.001, VP16: *p* > 0.05, two-way ANOVA of occupancy group and sgRNA). We also measured the radial BRD4 and H3K27ac enrichment profiles and observed a robust enrichment even with reduced VPR residence times (**Fig. 6G**). This corroborates our previous findings that BRD4 recruitment and histone acetylation occur efficiently even at high AD turnover rate. The residence time dependency of activation shown in **Fig. 6F** suggests that the TF is involved in an energy-dependent activation step. It is illustrated in a kinetic model how the residence time of the TF in the bound state can become a key determinant of transcription output independent of binding site occupancy (**Fig. 6H, Supplemental Information**). Furthermore, the higher sensitivity of VP16 mediated activation to a reduction of its binding affinity and residence time suggests that the higher multivalency of VPR can at least partly compensate for a reduction of direct DNA binding affinity.

**Fig. 6.**
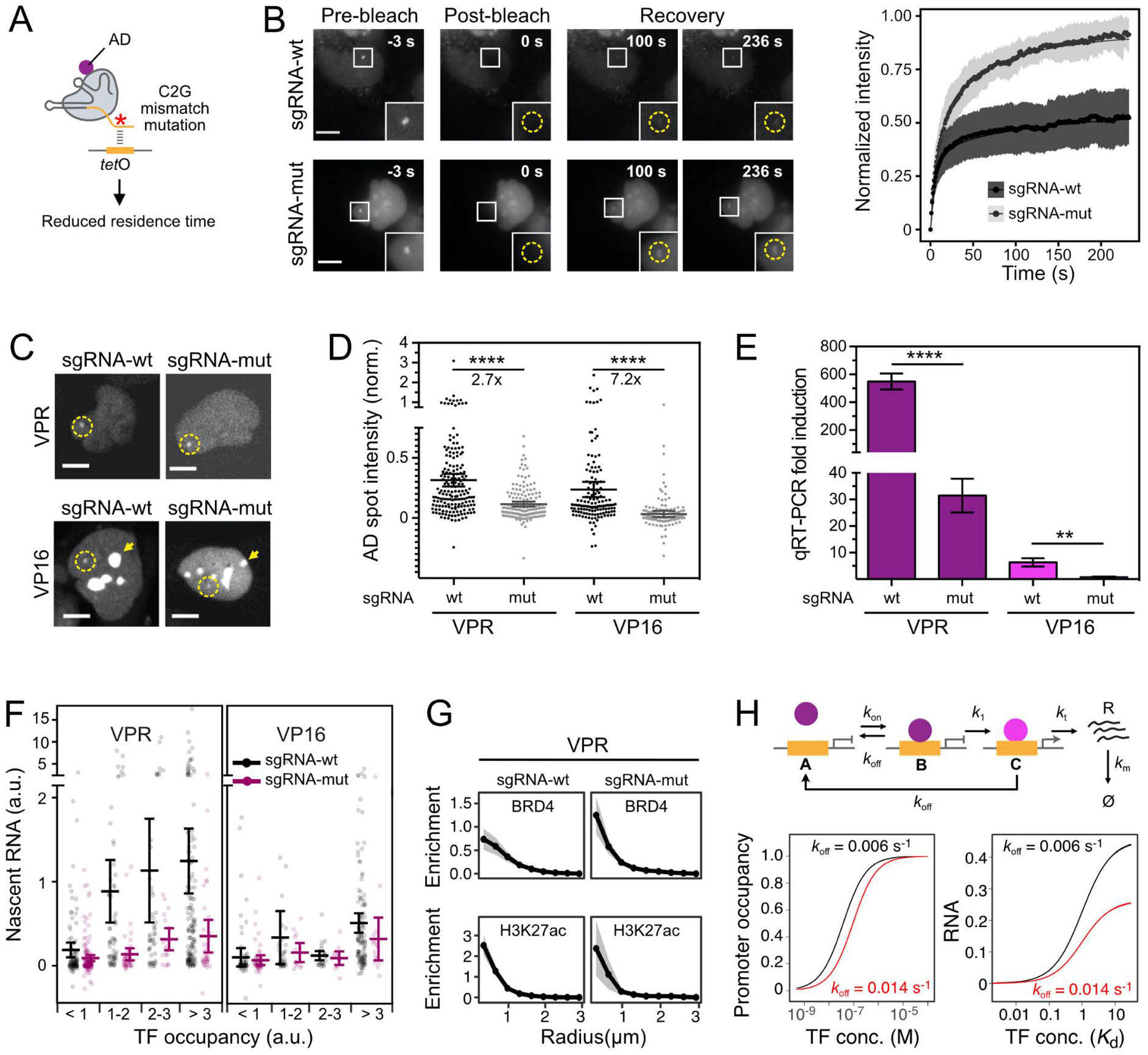
Effect of binding site occupancy, residence time and multivalent interactions on transcription activation. (**A**) dCas9 DNA binding affinity was lowered by introducing a C2G mismatch mutation in the targeting region of the sgRNA. (**B**) FRAP image series (left), averaged recovery curves and 95% CI (right) of dCas9-GFP-VPR and *tet*O sgRNA-wt (*n* = 10) or sgRNA-mut (*n* = 7). (**C**) Microscopy images showing enrichment of dCas9-GFP-VP16/-VPR at binding sites for sgRNA-wt/mut. Note that the VP16 construct is strongly enriched in nucleoli (arrows). The position of the reporter gene array is identified in the tagBFP-LacI marker channel (not shown, dashed circle). (**D**) Enrichment of dCas9 AD fusions with wildtype and mutated sgRNA at the reporter array (*n* = 127-175 cells per condition). Intensities were background subtracted and normalized to tagBFP-LacI marker intensity. Solid bar, mean; error bars, 95% CI; **** *p* < 0.0001 from two-sided Welch’s *t*-test. (**E**) qPCR measurements of reporter RNA levels 24 h after transfection for sgRNA-wt/mut. Mean and s. d. of fold-change reporter RNA induction compared to mock transfected samples and normalized to beta actin mRNA are shown (*n* = 3). The data for sgRNA-wt displayed in **Fig. 5A** is included for comparison. **, *p* < 0.01; ****, *p* < 0.0001 from two-sided unpaired Student’s *t*-test. (**F**) Nascent RNA detected via tdMCP-tdTomato 24 hours after transfection normalized to tagBFP-LacI marker intensity. Cells were divided into groups with equal TF occupancies determined from the GFP-AD signal normalized to the marker. Transparent dots correspond to values for single cells; mean and 95% CI are indicated. Note the axis break to visualize the majority of cells as well as the few cells with very high RNA production in the same plot. (**G**) Qualitative detection of activation marks by radial enrichment profiles of mCherry-BRD4 and H3K27ac immunostaining for dCas9-GFP-VPR. Data for sgRNA-wt from **Fig. 5 D** is shown for comparison. Mean and 95 % CI; *n* = 28-184 cells per condition. (**H**) Model for a multi-step transcription activation mechanism showing the dependence of RNA production at saturated binding on TF residence time (**Supplemental Information**). After TF binding to the promoter (state *B*) induction of transcription requires another energy consuming transition to state *C* (indicated by color change) where RNA is produced from the TF-bound promoter with rate *k*_1_. Two different dissociation rates *k*_off_ = 0.006 s^-1^ (*τ*_res_ = 167 s) and *k*_off_ = 0.014 s^-1^ (*τ*_res_ = 71 s) were compared. Binding site occupancy was computed for *k*_on_ = 10^5^ M^-1^ s^-1^, corresponding to a *K*_d_ of 60 nM and 140 nM, respectively. Steady state RNA levels computed for the two *k*_off_ values are shown as a function of TF concentration in *K*_d_ units, i. e. for the same promoter occupancy.

## Discussion

We combined different DBD and AD modules into a panel of synthetic TFs to dissect contributions of DNA binding properties, multivalent AD interactions, phase separation, co-activator recruitment and histone acetylation to transcription activation. The main results are summarized in the scheme depicted in **Fig. 7**. We show that TF activation strength is reduced with shorter residence time independent of TF binding site occupancy (**Fig. 6F**). This finding corroborates conclusions from previous studies (Brouwer and Lenstra, 2019; Callegari et al., 2019; Clauss et al., 2017; Gurdon et al., 2020; Loffreda et al., 2017; Shelansky and Boeger, 2020). It suggests that TF binding is coupled to an energy dependent kinetic proofreading step like nucleosome remodeling (Shelansky and Boeger, 2020), promoter DNA melting (Osman and Cramer, 2020) or TF post-translational modification (Qian et al., 2020). The activation strength of the five ADs studied here correlated with their capacity to engage in multivalent interactions as assessed by their propensity to form optodroplets (**Fig. 2A-D**). The comparison of two prototypic ADs with high (VPR) versus low (VP16) multivalency provided insight into the underlying molecular differences. The analysis of TF chromatin binding by FRAP and the experiments that employed a mutated sgRNA revealed that multivalent interactions stabilized chromatin binding and led to the accumulation of additional TF molecules via protein-protein interactions at the promoter (**Fig. 4C, D**). Furthermore, recruitment of co-activators like BRD4 and setting the H3K27ac modification was clearly enhanced by VPR as compared to VP16 (**Fig. 5D-I**). Thus, we conclude that multivalent TF interactions foster interaction both with the TF itself but also with co-factors such as BRD4 or p300, thereby boosting transcription or increasing persistence of the activated state. This is in line with previous reports on the ability of BRD4 to engage in dynamic multivalent interactions (Cho et al., 2018; Han et al., 2020; Ma et al., 2021; Sabari et al., 2018). It is noted that BRD4 was neither required nor sufficient to induce transcription in our experiments (**Fig. 5D, S5D-E**). Furthermore, transcription was not a prerequisite for BRD4 binding or H3K27 acetylation (**Fig. 5D**). Interestingly, BRD4 enrichment around the promoter could also be maintained upon weakened DNA binding in the sgRNA mutation experiment **(Fig. 6 G)**. We conclude that maximum transcription activation strength requires the stable binding of the AD at the promoter with long residence times for interactions with the core transcriptional machinery, while co-activators can also be recruited to the promoter by TFs at high turnover rate.

**Figure 7.**
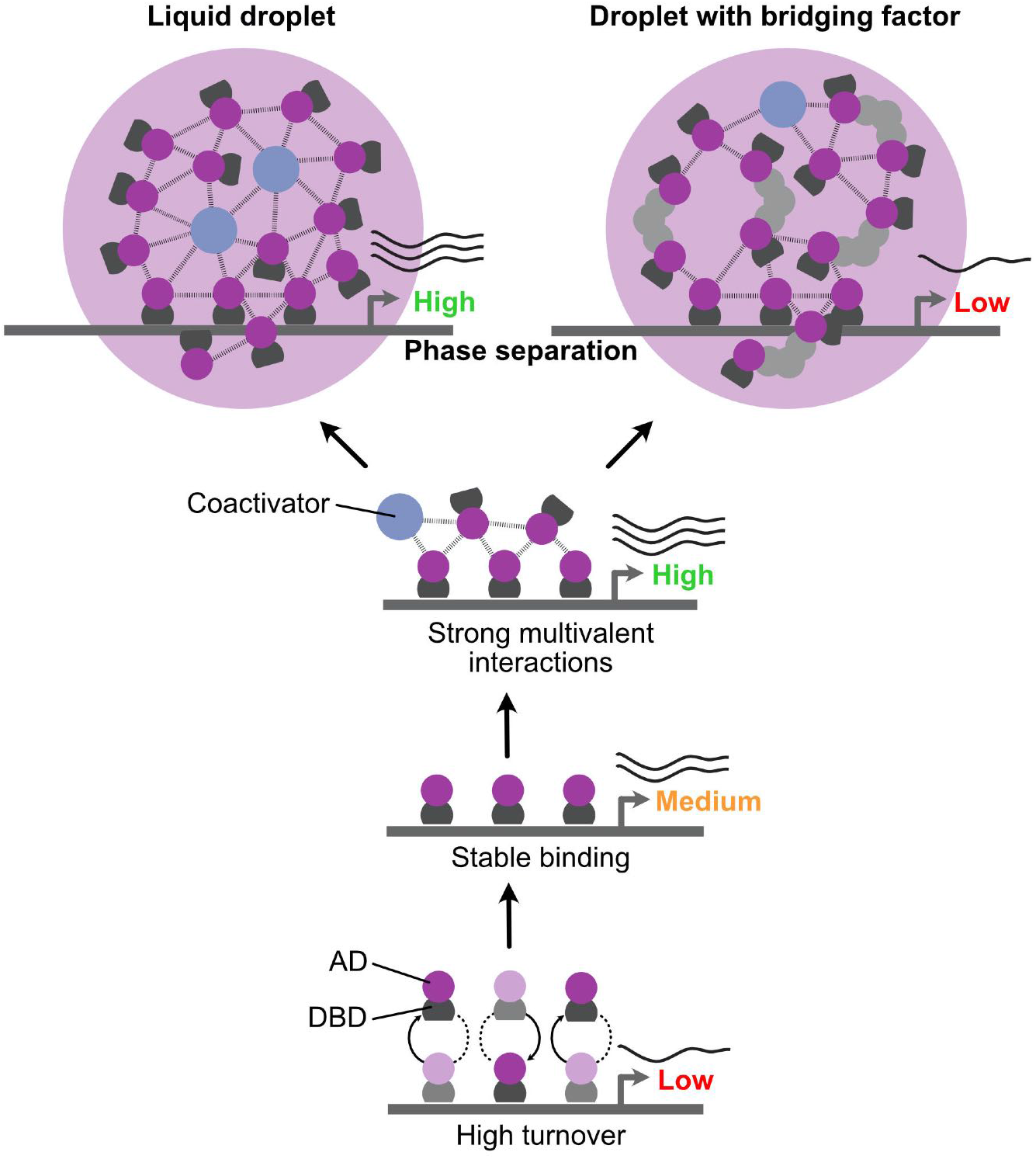
Model for dependence of transcription on TF promoter occupancy, residence time and multivalent interactions. From bottom to top: Low residence times lead to lower transcription (high turnover vs. stable binding) and multivalent AD interactions increase transcription activation capacity, in part via interactions with coactivators. The formation of phase-separated droplets can further increase the local TF concentration but does not further increase RNA production (top left). Rather, a multimeric TF assembly stabilized by introducing additional interactions via bridging factors can create a repressive subcompartment despite a strong TF enrichment at the promoter (top right).

Recent studies investigated the functional consequences of TF phase separation and reported that droplets formed by light induction of PHR/CRY2-TAF15 (Wei et al., 2020) or IDR-VP16 constructs (Schneider et al., 2021) amplify gene expression or increase transcription activation. Furthermore, it has been proposed that phase separated transcriptional condensates that include super-enhancers drive transcription of highly active genes (Hnisz et al., 2017; Sabari et al., 2018; Shrinivas et al., 2019). Our results confirm an enhancement of transcription when fusing the N-terminal IDR of FUS to the activation domain VP16 in our light-inducible TF setup (**Fig. 3G-I**). FUSN-VP16 also behaved differently from the VP16 droplet state enforced by bridging factors, suggesting that the intrinsically disordered FUSN domain provides interactions that favor efficient transcription initiation. Previous studies attributed this effect to the local TF enrichment by LLPS from the comparison to control constructs without IDRs and/or in the absence of a light trigger. However, experiments that assess transcription activity of the same chromatin bound activator in the presence/absence of droplet formation under identical conditions were lacking. Here, we demonstrate that it is not justified to generally assume that liquid TF droplets at the promoter would enhance transcription as we see a neutral effect when comparing activation both below and above the critical droplet forming concentration for the same activator (**Fig. 2E-G**). Based on these findings, we argue that assemblies formed by weak multivalent interactions as established by the VPR or FUSN domain are sufficient to enhance transcription without phase separation. Instead, we propose that phase separation of TFs at artificially high protein concentration simply reflects a physico-chemical property of IDRs in line with previous finding (Chong et al., 2018). As shown here, stabilization of TF binding at the promoter and TF dependent co-factor recruitment are two possible mechanisms by which multivalent interactions could enhance transcription in the absence of phase separation. In the experiments that induced TF droplets via the addition of bridging factor we observed an inhibitory effect on transcription (**Fig. 3**). Interestingly, a conceptionally related mechanism of transcription suppression has been observed for the sequestering of RNA polymerase I into a phase-separated nucleolar cap subcompartment (Ide et al., 2020). Whether inhibitory phase-separated states are involved in regulating Pol II activity in an endogenous cellular environment, for example to establish refractory promoter states (Rodriguez and Larson, 2020), remains to be demonstrated.

In summary, our study reveals how DBD and AD activities are linked and jointly affect TF binding and activation properties. It provides novel insight into how transitions between active and inactive promoter states can be regulated. Furthermore, we anticipate that our findings are helpful for the design of CRISPR/dCas9-based synthetic TFs for control of gene expression programs (Pandelakis et al., 2020), and will stimulate further experiments that dissect how phase-separation processes affect Pol II transcription.

## Author contributions

Study design: JT, LF, KR. Acquisition of data: JT, LF, AR, PG. Analysis of data: all authors; Drafting of manuscript: JT, LF, KR. Manuscript reviewing: all authors. Supervision and coordination: KR

## Acknowledgements

We thank Robin Weinmann and Fabian Erdel for discussion and the DKFZ light microscopy core facility for technical support. This work was funded by Priority Program 2191 “Molecular Mechanisms of Functional Phase Separation” of the Deutsche Forschungsgemeinschaft (DFG) by DFG grant RI1283/16-1 to KR. Data storage at SDS@hd was supported by the Ministry of Science, Research and the Arts Baden-Württemberg (MWK) and the DFG through grants INST 35/1314-1 FUGG and INST 35/1503-1 FUGG.

## Materials and Methods

### Plasmids

Plasmids and sgRNA sequences are listed in **Supplemental Table S1 and S2**. Protein constructs were expressed under control of a CMV promoter using pEGFP-C1/N1 (Clontech) (enhanced GFP, referred to here as GFP) or pcDNA3.1 (Invitrogen) vector backbones. Plasmids expressing *lacO*/*tetO* targeting sgRNAs with 2xPP7 loops were designed as gBlocks (Integrated DNA Technology) and cloned into a U6 promoter-driven sgRNA expression vector derived from Addgene plasmid #61424, and with the PP7 loop sequence reported previously (Zalatan et al., 2015). The dCas9 open reading frame used in all dCas9-based constructs originates from Addgene plasmid #60910. Activation domains were obtained from Addgene plasmid #63798 (VPR, p65, Rta) (Chavez et al., 2015), Addgene plasmid #103836 (VP16) (Gunther et al., 2013) and pSTAT2-EGFP (STAT2, amino acids 1-10 and 722-857) (Frahm et al., 2006). The p300 core histone acetyltransferase domain was derived from Addgene plasmid #61357. CRY2PHR and CIBN domains were taken from Addgene plasmid #26866 and #26867, respectively. The CIBN-dCas9-CIBN expression construct corresponds to Addgene plasmid #60553. Tandem MCP with tandem Tomato (tdMCP-tdTomato) was derived from Addgene plasmids #40649 and #54642. The TATA-box of the promoter was removed to reduce expression levels. The tandem PCP (tdPCP) protein was derived from Addgene plasmid #40650. LacI and TetR constructs are based on the previously described fluorescently tagged proteins (Lau et al., 2003)(Pankert et al., 2017). The rTetR protein sequence was subcloned from the Tet-On transactivator used in the commercially available Tet-On 3G system (Takara Bio). FUSN was derived from Addgene #122148 and GBP was based on a previously described construct (Rothbauer et al., 2008).

### Cell Culture

U2OS 2-6-3 cells containing the stably integrated *lacO/tetO* reporter gene cluster (Janicki et al., 2004) were grown in DMEM (1 g/l glucose, Gibco) without phenol-red supplemented with 10 % tetracycline-free fetal calf serum (FCS), penicillin/streptomycin and 2 mM L-glutamine using standard cell culture methods at 37 °C and 5 % CO_2_. Cells were seeded onto 8-well chambered coverglass slides (Nunc Labtek, Thermo Fisher Scientific) at a density of 2·10^4^ cells per well. For qPCR 3·10^5^ cells were seeded in 6-well plates. One day after seeding, the medium was replaced with imaging medium (FluoroBrite, Gibco, A1896701; 10 % tet-free FCS; penicillin/streptomycin; 2 mM L-glutamine) and cells were transfected using the Xtreme-Gene 9 reagent (Roche) according to the manufacturer’s guidelines. Briefly, 200-400 ng plasmid DNA and 0.6 µl transfection reagent in 20 µl OptiMem (Gibco) were used per well for microscopy experiments. The plasmid DNA mix consisted of 100 ng of guide RNA plasmid and 100 ng of equal amounts of the remaining constructs. For transfections without guide RNA plasmid the 200 ng were split equally among the plasmids. Transfection reactions were scaled up to 2 µg plasmid DNA per well for qPCR experiments. Cells were protected from light until the start of experiments for FRAP and induction time course experiments with light-responsive constructs. FRAP experiments were conducted 48 hours post-transfection, all the other experiments 24 hours post-transfection. For the radial profile microscopy experiments or qPCR of light-inducible activator constructs, cells were illuminated by diffuse white LED light for 24 hours. rTetR activator constructs were allowed to bind in presence of 5 µg/mL doxycycline (Sigma-Aldrich, D9891) which was added after transfection.

### RNA isolation and qPCR

Total RNA was isolated using QIAzol lysis reagent (Qiagen), followed by one round of chloroform extraction and isopropanol precipitation. The purified RNA was treated for 30 min at 37°C with RQ1 DNase (Promega) according to the manufacturer’s protocol and then purified using one round of each phenol/chloroform and chloroform extraction followed by precipitation using ethanol in presence of 300 mM sodium acetate pH 5.5 and GlycoBlue coprecipitant (Thermo Fisher Scientific). RNA concentration and purity were determined by absorbance measurement. Per sample, one microgram of DNase-treated RNA was used as input for cDNA synthesis using the Superscript IV reverse transcriptase protocol (Thermo Fisher Scientific). qPCR was carried out in technical triplicates with 2 µl of 1:40-diluted cDNA per 10 µl reaction using SYBR Green PCR Mastermix (Applied Biosystems) with a final primer concentration of 500 nM. The following PCR primers (Eurofins Genomics) were used. Human beta-actin fwd: 5’-TCC CTG GAG AAG AGC TAC GA-3’, rev: 5’-AGC ACT GTG TTG GCG TAC AG-3’; VPR-VP16 fwd: 5’-AAGAAGAGGAAGGTTGCCCC-3’, rev: 5’-CCC CAG GCT GAC ATC GGT-3’; CFP-SKL fwd: 5’-GTCCGGACTCAGATCTCGA-3’, rev: 5’-TTC AAA GCT TGG ACT GCA GG-3’. The qPCR analysis was carried out using the 2^-ΔΔCT^ method. Reporter RNA expression levels (CFP-SKL) were normalized to beta-actin mRNA levels (ΔCT) and then expressed as fold-change of the mock control.

### Microscopy instrumentation

SRRF images and data for radial profiles and occupancy were acquired with an Andor Dragonfly 505 spinning disc microscope equipped with the Nikon Ti2-E inverted microscope and a 40x oil immersion objective (CFI Plan-Fluor 40x Oil 1.30/0.20, Nikon). Multicolor images were acquired using laser lines at 405 nm (tagBFP), 488 nm (GFP), 561 nm (tdTomato and mCherry) and 637 nm (Alexa 633) for excitation with a quad-band dichroic unit (405, 488, 561, 640 nm) and corresponding emission filters of 450/50 (tagBFP), 525/50 (GFP), 600/50 (tdTomato, mCherry) and 700/75 nm (Alexa 633) and an iXon Ultra 888 EM-CCD camera. Live cell experiments were conducted in an incubation chamber (Okolab) at 5 % CO_2_ and 37 °C temperature. Light-induced time course and FRAP experiments were carried out with an AxioObserver Z1 widefield microscope (Zeiss) equipped with a 20x air objective (Zeiss Plan-Apochromat 20x/0.8 M27), the Zen 2012 pro software including modules for z-stack, time-lapse and multi-position acquisition and an AxioCam MRm Rev.3 monochrome camera with filter sets with excitation bandpass, beam splitter, emission bandpass wavelength: GFP, 470/40 nm, 495 nm, 525/50 nm; tdTomato, 535/30 nm, 570 nm, 572/25 nm and mCherry, 550/25 nm, 590 nm, 629/62 nm. For spot bleaching at 473 nm in FRAP experiments the microscope was extended with an UGA40 70 mW laser scanning system (Rapp OptoElectronic). A Leica TCS SP5 II confocal microscope (Leica) equipped with a 63x Plan-Apochromat oil immersion objective was used for additional FRAP experiments as described previously (Muller-Ott et al., 2014) and for measuring recruitment/dissociation kinetics (**Supplemental Information**).

### Super-resolution radial fluctuations (SRRF) imaging of reporter locus

Cells were transfected with LacI containing a SNAP-tag and the respective components of the activation complex directed to the *tetO* sites. After 24 hours SNAP-Cell 647-SiR substrate (New England Biolabs) was added to the medium at a concentration of 3 μM, incubated for 30 min and washed three times with medium, incubated with medium for 30 min and washed three times with PBS. Cells were fixed with 4 % paraformaldehyde for 10 min and washed with PBS before imaging. Imaging was performed on the Andor Dragonfly spinning disc microscope with a 100x silicon immersion objective (CFI SR HP Apochromat Lambda S 100x, Nikon) and a 2x magnification lens to ensure oversampling. 200 frames were acquired per channel for one super-resolved SRRF image. Exposure time was 2.5 ms with 100 % laser intensity of the 488 nm or 637 nm laser for GFP or 647-SiR, respectively. SRRF analysis was performed using the SRRF-stream tool implemented in the microscope software with 5 × 5 sub-pixels, a ring radius of 1.5 pixels for radiality calculations and mean-projection of radiality images.

### Light-induced time course experiments

Light-induced time course experiments followed the protocol given in (Trojanowski et al., 2019) and were conducted with the AxioObserver Z1 widefield microscope. Slides with transfected cells were kept in the dark until the start of image acquisition and red-light illumination was used during sample preparation before initiating the reaction with blue light. For JQ1 (Sigma-Aldrich, SML1524) treatment the drug was diluted in medium and added to the respective wells to a final concentration of 1 µM three hours before the start of imaging. Doxycycline (Sigma-Aldrich, D9891) was added 15 minutes before imaging to a final concentration of 5 µg/ml in the dark to induce binding of CIBN-rTetR. The focal plane was determined by red-filtered transmitted light and kept constant by the hardware autofocus. Imaging time courses comprised repeated cycles of imaging of a grid of 16 positions (4×4, 50 % negative overlap) with three z-slices (distance 1.0 µm) in intervals of 2 minutes over 90 minutes or 60 minutes for BRD4 recruitment experiments. After each time course experiment the slide was exchanged with a slide that had been stored in the dark to ensure that between experiments PHR molecules that had been exposed to stray light from a neighboring well had reverted to their inactive conformation in the dark.

### Analysis of time course images

Image were processed with the EBImage and NSSQ R packages as described previously (Trojanowski et al., 2019). In a first step positions of nuclei with successful recruitment of PHR-GFP-AD were manually selected and segmented in the GFP channel by automated local thresholding for each time point. The nucleus was tracked by mapping the segmented objects with minimal distance in consecutive frames. The best focal plane was selected from the z-stack for each time point using the intensity gradient inside the nucleus area. The reporter gene cluster was segmented inside the nuclear area using a quantile-based threshold in the PHR-GFP-AD channel. The spot position was tracked through the time course by finding the closest segmented object in consecutive images. The areas of spot (*A*_*spot*_) and nucleus masks (*A*_*nucleus*_) and the average intensities inside the spot (*I*_*spot*_), nucleus (*I*_*nucleus*_) and ring-shaped background regions around them (*I*_*spotbg*_, *I*_*nucleusbg*_) were measured in each channel. The amount of fluorescence intensity recruited to the reporter spot was then calculated as the product of background subtracted spot intensity and area:

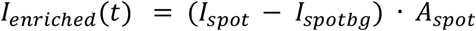

Segmented image series were manually curated by removing cells with morphological abnormalities, missing expression or segmentation errors and then classified as responders or non-responders based on visible accumulation of intensity in the reporter spot in the reader channel. In order to account for the small time shift of acquisition between positions in one imaging cycle, the intensity values at the beginning of each cycle were calculated by linear interpolation. The resulting single cell time courses were then either directly averaged for each time point to yield absolute intensity values or normalized by subtracting the initial value and dividing by the maximum value before averaging:

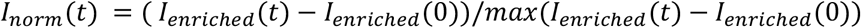

Averaging was performed either for all cells or only for responder cells. The first value was subtracted so that all curves started at an intensity value of zero. For BRD4 recruitment time courses time traces were normalized to their maximum values without subtraction of the first time point. The value of the first time point was then subtracted after averaging. Times to half-activation were determined from single cell time courses as the first time point, at which the normalized intensity equaled or exceeded 0.5. The responder fraction was calculated as the number of cells annotated as responders divided by the total number of cells remaining after manual curation. Time course maximum values were determined as the average plateau value of tdMCP-tdTomato intensity over the last five time points.

### Light induced optodroplet formation

Image series of cells transfected with combinations of PHR-GFP-AD and CIBN-rTetR, CIBN-dCas9-CIBN or dCas9 + tdPCP-CIBN were acquired in the GFP channel with the same settings as the induction time course experiments over 6 cycles at 25 positions. For conditions with CIBN-rTetR doxycycline was added 15 minutes before imaging. All images were acquired in a single microscopy session and processed as described above. To remove the contribution of the reporter array spot to the nuclear intensity variation, the reporter spot was selected manually and removed from the nucleus mask using a disc shaped area with 7 pixels diameter. Mean and standard deviation of intensities in the processed nucleus images and in a ring-shaped background area around the nucleus were determined. Subsequently, image series were manually curated and classified as containing optodroplets or not by checking for the presence of spherical structures outside the reporter spot. For quantification, optodroplets inside the nucleus were segmented using the median of the nuclear intensity multiplied by 1.75 as a segmentation threshold. The droplet abundance was determined as the area of segmented droplets normalized by the nucleus area. The critical value for droplet formation was determined as the nuclear intensity at which the relative droplet area exceeded an empirically determined threshold of 1%. This threshold yielded a good agreement with the manual annotation of cells as droplet containing or not. In order to represent the relative droplet area as a smooth function of the nuclear intensity we fitted a logistic function to it:

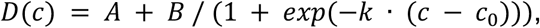

where *D(c)* is the droplet abundance, *c* the nuclear concentration and the remaining free fitting parameters are offset *A* and amplitude *B*. The intensity at which this function crosses 1% corresponds to the critical value.

### FRAP analysis

FRAP experiments were carried out on the Zeiss widefield microscope described above with an external micromanipulation laser for bleaching. This set-up allowed fast acquisition of time courses in a large number of cells and conditions and yielded results similar to those obtained with a confocal microscope (**Fig. S4 A-C, Supplemental Information**). Laser position calibration was performed according to the UGA40 software instructions on a fluorescent calibration slide. Conditions with optogenetic constructs were illuminated for at least one minute in the GFP channel to saturate binding to CIBN before carrying out FRAP. The reporter spot was manually selected as bleach region and bleached at 100 % laser intensity for one second, 3-4 frames after starting an imaging time series of four minutes with one second intervals (on-spot bleach). For determining construct-specific diffusion coefficients a central nuclear bleach region outside of the reporter spot was bleached and fluorescence recovery was monitored at 300 ms intervals for one minute (off-spot bleach). FRAP of *tetO*-bound dCas9-GFP-VPR with sgRNA-wt and sgRNA-mut was carried out with the same settings but with an alpha Plan-Apochromat 100x/1.46 Oil DIC M27 objective (Zeiss). The spot or bleach region intensities in the image series were quantified by a semi-automated analysis pipeline with our *R* software package *NSQFRAP* and normalized to pre-bleach and nuclear intensity to account for bulk bleaching. Normalized recovery curves were fitted with a reaction-diffusion model for clustered binding sites (Sprague et al., 2006) using the empirical post-bleach profile as an initial condition and the effective diffusion coefficient determined from the off-spot FRAP measurements (**Fig. S4D, Supplemental Information**).

### Analysis of binding and dissociation kinetics

Binding and dissociation time courses of PHR-mCherry-VP16 on CIBN constructs were performed on a Leica SP5 confocal microscope. Transfected cells were imaged using a 594 nm laser line (mCherry) for focusing without triggering the optogenetic components. An image was taken before starting a 2-3 min image series using both the 594 nm and 488 nm laser line for imaging the co-transfected GFP-LacI array marker and triggering PHR-CIBN interaction with time intervals of 6 seconds. After recoding this time series, the 488 nm laser line was switched off and 2 µm z-stacks (0.5 µm step-size) of the same positions were recorded for 20-30 min at 1 min intervals to monitor PHR-CIBN dissociation. Reporter spot tracking and intensity quantification were performed as described for FRAP and using the GFP-LacI array marker to identify the reporter spot. Spot intensities were subtracted from the background intensity determined in a ring-shaped area around the spot and normalized for each cell to the last (*t* = 168 s, binding) or first timepoint (*t* = 0 s, dissociation), respectively.

### Immunofluorescence

Slide wells with transfected cells that had been illuminated for 24 hours were washed once with phosphate-buffered saline (PBS) and fixed with 4 % paraformaldehyde in PBS (Sigma-Aldrich, 252549) for 12 minutes. After washing with PBS cells were permeabilized with ice-cold 0.1 Triton-X100 (Merck, 108643) in PBS for 5 minutes. Blocking with 10 % goat serum (Cell Signaling Technology) in PBS for 15 minutes was followed by incubation with rabbit anti H3K27ac antibody (ab4729, Abcam, Lot GR183922-1, 1:1000) in 10 % goat serum for one hour at room temperature. Cells were washed three times for 5 min with 0.002 % NP40 (Sigma-Aldrich, i8896) in PBS. Incubation with the secondary antibody goat anti-rabbit Alexa 633 (Thermo Fisher Scientific, A21071, Lot 1073053, 1:1000) was done for 30 minutes at room temperature in 10 % goat serum/PBS. Cells were washed twice for 5 min with PBS and stored in PBS at 4 °C until they were imaged on the following day.

### Single molecule RNA FISH

Probes of the RNAScope system (ACD Bio) against the MS2 sequence of the U2OS-2-6-3 reporter cell line covering position 851 to 2163 of the reporter RNA were custom designed by ACD Bio. Slides with transfected cells that had been illuminated for 24 hours were washed once with phosphate-buffered saline (PBS) and fixed with 4 % paraformaldehyde in PBS (Sigma-Aldrich, 252549) for 12 minutes. After washing three times with PBS cells were treated with 3 % hydrogen peroxide for five minutes, washed with PBS and treated with protease III (ACD Bio) diluted 1:15 for ten minutes, followed by three PBS washes. Hybridization of target and amplification probes was then performed according to the manufacturer’s protocol. Target probes after amplification were labelled with Alexa Fluor 488 using the C1 detection kit. Cells were stored and imaged in PBS.

### Spinning disc confocal microscopy for radial profiles and occupancy plots

Cells transfected with light-responsive constructs were cultured for 24 hours in the presence of diffuse white LED light after transfection and then subjected to imaging. For each condition 14 µm z-scans (1 µm step size) on at least 81 positions (9 × 9 grid, 1 % overlap) were recorded per slide well on an Andor Dragonfly 505 spinning disc microscope. Images were processed with the *NSSQ* package (Trojanowski et al., 2019). Nuclei with activator recruitment in the GFP-AD channel or array marker signal were manually selected in maximum projections of each position and then segmented in sum-projected images by local thresholding. Three consecutive z-planes with the highest contrast were mean-projected to yield a single image for quantification. Subsequently, the spot position was selected in each segmented cell based on the co-transfected array marker (tagBFP-LacI) and a disc shaped spot mask with a diameter of 1.6 µm (5 pixels), a ring-shaped background mask and a nuclear mask were used to quantify average intensities in all channels. Spot mask diameters were 1.6 µm (5 pixels) for activators and 3.8 µm (12 pixels) for nascent RNA. Radial profiles were measured by creating masks of concentric rings of pixel-wise increasing radius around the spot position and measuring average intensities up to a radius of 2.9 µm (9 pixels). The minimum value was subtracted from the profiles and they were divided by the local background intensity for normalization. Single cell profiles were averaged for each condition and the minimum value was subtracted. The resulting enrichment score profile gives qualitative information about the accumulated intensity in the spot center. For quantitative comparisons of local concentrations (occupancy and promoter activity plots), average spot intensities were measured in images acquired on the same day with the same imaging parameters. The average intensity in the spot background region was subtracted from the average spot intensity. The resulting intensity in the activator-GFP channel was normalized to the tagBFP-LacI marker channel.

### Statistics, data presentation and analysis software

Mean values and 95 % confidence intervals (CI) for time courses of nascent transcripts, BRD4 or fluorescence recovery and for intensity profiles were calculated from single cell data for every time point or radial position from a Student’s t-distribution. Pairwise comparisons of the mean for qPCR or relative intensities were done using unpaired, two-sided Student’s or Welch’s t-tests, respectively. To check for the respective effects of two grouping variables (AD type and presence of optodroplets for half-activation times; occupancy group and sgRNA for the effect of residence time on promoter activation) a two-way ANOVA (type II) was performed. Error bars represent one standard deviation (s. d.) for qPCR experiments and the standard error of the mean (s. e. m.) for half-activation times as indicated. For residence times the mean and CI of *k*_off_ were determined before calculating the inverse (1/*k*_off_). Axis breaks were introduced in relative intensity and qPCR plots for conditions with values on very different scales or with outliers and are marked by an interruption of the axis. Box plots show first and third quartile (box), median (bar), data points within 1.5-fold interquartile range (whiskers) and outliers (points). Images were processed with the *NSSQ* package (Trojanowski et al., 2019) available at https://github.com/RippeLab/NSSQ. Exemplary microscopy images were linearly adjusted for visibility using Fiji with the same adjustments applied for all time points (Schindelin et al., 2012). The *R* software package *NSQFRAP* for the semi-automated FRAP analysis can be downloaded from https://github.com/RippeLab/NSQFRAP.

## Notes

### Competing Interest Statement

The authors have declared no competing interest.

