## Supplemental Information for "Transcription activation is enhanced by multivalent interactions independent of phase separation"

### Content

#### Supplemental methods

FRAP data analysis
Model for TF residence time dependent activation
Characterization of transcription incompetent complexes

#### Supplemental figures

Figure S1. Light induced activator binding and transcription activation
Figure S2. AD droplet formation propensity and transcription activation kinetics
Figure S3. Effect of droplet formation on transcription activation by VP16
Figure S4. Experimental FRAP setup and data analysis
Figure S5. Histone acetylation, BRD4 binding and transcription activation
Figure S6. Modulation of TF residence time via sgRNA mutations

#### Supplemental tables

Table S1. Plasmid constructs
Table S2. sgRNAs sequences used for dCas9 targeting
Table S3. Propensity of the activation domain to form optodroplets
Table S4. Transcription activation kinetics
Table S5. FRAP parameters of TF dynamics
Table S6. Reporter RNA expression measured by qRT-PCR
Table S7. Histone acetylation, BRD4 binding and transcription activation
Table S8. Binding site occupancy of dCas9

### Supplemental analysis

### FRAP analysis

#### Comparison of FRAP for widefield and confocal microscopy setup

The widefield microscope FRAP used in our study provides fast data acquisition and imaging is decoupled from bleaching. However, the resolution along the z-axis is not as good as with a confocal microscopy. We thus compared GFP-LacI diffusion and binding to the reporter array in FRAP experiments with the widefield FRAP system to measurements with a Leica SP5 confocal microscope (Leica, Germany) equipped with a 63x Plan-Apochromat immersion objective. The confocal FRAP experiments were conducted using the Leica LAF software and bleaching with the argon laser lines (458 nm, 476 nm, 488 nm, 496 nm). Images of 128x128 pixel with zoom factor 9 corresponding to 194 nm/pixel were recorded at 1400 Hz line frequency resulting in a frame time of 115 ms. For each cell, 70 pre-bleach and 2 bleach frames were recorded with a 1 µm diameter circular bleach region. It was placed on the reporter array ("on spot") or elsewhere in the nucleus but outside nucleoli ("off spot"). Subsequently, 1200 (on spot) or 300 (off spot) post-bleach frames were recorded (**Fig. S4A-C**). Image analysis and parameter estimation were done as for widefield FRAP with the following adaptations: The post-bleach intensity profiles were not fitted individually but averaged, the estimated bleach profile parameters were applied globally to calculate initial conditions and the fitting range of the correction factor *bgRatio* was set to 0.3 to 2.3. A fit of the data to reaction-diffusion model yielded similar values for the two different FRAP setups of $D_{eff}$ = 2.3 µm$^2$/s and $k_{off}$ = 0.009 s$^{-1}$ (widefield) vs. $D_{eff}$ = 3.3 µm$^2$/s and $k_{off}$ = 0.010 s$^{-1}$ (confocal) (**Fig. S4B, C**). The widefield curves recovered to higher values in the first seconds and then showed a similar behavior as the confocal FRAP curves but with a lower immobile fraction (widefield: 7.8 %, confocal: 29 %). These differences can be rationalized by the better z-resolution of the confocal setup that reduces the number of freely diffusive molecules observed below and above the reporter array, which do not contribute to a potential immobile fraction. Moreover, shorter FRAP time courses were recorded with the confocal system due to higher imaging related bleaching and out-of-focus translocation of the reporter array. Potentially, this shorter observation time in confocal mode may lead to a higher estimate of the immobile fraction.

#### Analysis of FRAP images

Intensities in the region of interest were determined automatically using functions of the NSSQ (Trojanowski et al., 2019) and EBImage (Pau et al., 2010) packages in R (R Core Team, 2020) and the bleached nucleus was segmented by local thresholding (**Fig. S4D**). As dCas9-GFP was depleted in the nucleus, images were blurred and the whole cell was segmented for this construct. For on-spot experiments the reporter array was segmented in the first pre-bleach frame using the 98% quantile inside the nucleus. The bleach region was segmented in an image created from the difference of pre-bleach and first post-bleach frame. To correct for chromatin or cell movements the nucleus was tracked, and positions of spot and bleach region mask were shifted accordingly. If automated tracking failed, spot positions were selected manually in every tenth frame and all masks were shifted accordingly. Average intensities were extracted for each time frame in the nucleus, in a ring-shaped area around the nucleus (background intensity), in the spot area and in a ring-shaped area around the spot (local background). The intensity profile around the center of the bleach position was measured as the median intensity of rings starting with a radius of 1 pixel up to a radius of 9 pixels (20x objective) or 40 pixels (100x objective). The pixel size was 0.63 µm (20x objective) or 0.13 µm (100x objective) based on a reflective grid slide of known size. Recovery curves of profiles and average intensities were subjected to the following normalizations: Background $I_{nuc\_bg}$ in a region around the nucleus was subtracted and intensity profiles $I(r,t)$ were normalized to the average nuclear intensity $I_{nuc}$ to account for the overall reduction of fluorescence signal during the experiment. The intensity of the center position of the first post-bleach frame $I(r{=}r_{center}, t{=}0)$ was subtracted. The resulting profile was normalized to the average value before bleaching for each profile position $r$.

$$I_1(r,t) = I(r,t) - I_{nuc\_bg}(t)$$

$$I_{nuc\_norm}(t) = I_{nuc}(t) - I_{nuc\_bg}(t)$$

$$I_2(r,t) = \frac{I_1(r,t)}{I_{nuc\_norm}(t)}$$

$$I_3(r,t) = I_2(r,t) - I_2(r = r_{center}, t = 0)$$

$$I_{norm}(r,t) = \frac{I_3(r,t)}{mean\big(I_3(r, t < 0)\big)}$$

For off-spot experiments the average bleach region intensity was calculated from normalized profiles by averaging intensities from the region center to a radius of 3.5 µm weighted by the pixel number in each ring of the profile and leaving out the innermost value. For quantitating the spot intensity, the nuclear background signal was subtracted. Average spot intensities were normalized by dividing them by the average nucleus intensity, subtracting the minimum value in the first post-bleach frame and dividing by the average pre-bleach value.

$$I_{spot,1}(t) = I_{spot}(t) - I_{nuc\_bg}(t)$$

$$I_{nuc\_norm}(t) = I_{nuc}(t) - I_{nuc\_bg}(t)$$

$$I_{spot,2}(t) = \frac{I_{spot,1}(t)}{I_{nuc\_norm}(t)}$$

$$I_{spot,3}(t) = I_{spot,2}(t) - min\left(I_{spot,2}(t)\right)$$

$$I_{spot,norm}(t) = \frac{I_{spot,3}(t)}{mean\left(I_{spot,3}(t < 0)\right)}$$

Segmented image series were manually curated by removing cells where (i) segmentation or tracking failed, (ii) the normalized spot intensity exceeded 1.2, (iii) the spot intensity was less than 25% above background, or (iv) the recovery curve displayed strong intensity jumps.

**Models for clustered binding sites and diffusion**

Recovery of fluorescence intensity inside the spot area was modeled by a localized cluster of binding sites *b* inside a cylindrical volume of radius $r_s$ that can be bound by freely diffusing particles *f* to form a complex *c* according to the theoretical framework established previously (Sprague et al., 2006):

$$for\, r \le r_s:$$

$$\frac{\partial f(r,t)}{\partial t} = D_{eff} \cdot \nabla_r^2 f(r,t) - k_{on}^* \cdot f(r,t) + k_{off} \cdot c(r,t)$$

$$\frac{\partial c(r,t)}{\partial t} = k_{on}^* \cdot f(r,t) - k_{off} \cdot c(r,t)$$

$$for\, r > r_s:$$

$$\frac{\partial f(r,t)}{\partial t} = D_{eff} \cdot \nabla_r^2 f(r,t)$$

$$c = 0$$

Here, $D_{eff}$ is the effective diffusion coefficient that includes free diffusion and transient non-specific binding to chromatin. The apparent rate $k_{on}^*$ for binding to cluster sites includes the equilibrium concentration of free cluster binding sites. We extended this description by using a bleach region that can be larger than the spot area and modeled the initial conditions by a Gaussian function with a central plateau. It accounts for diffusion during the time between bleaching and the first post-bleach frame.

$$I(r < r_p, t = 0) = 0$$

$$I(r \ge r_p, t = 0) = A \cdot \left(1 - e^{\frac{-(r-r_p)^2}{\sigma}}\right)$$

In this equation, $r_p$ is the plateau radius, *A* the intensity of the unbleached peripheral region and $\sigma$ describes the width of the Gaussian. The binding site cluster was approximated as a cylinder with a homogeneous distribution of binding sites in z-direction at the center of a cylindrically shaped nucleus. This allowed us to formulate the system of partial differential equations in polar coordinates. For the estimation of diffusion coefficients from off-spot FRAP experiments we used a simplified model with only a diffusive and an unspecific immobile fraction.

$$\frac{\partial f(r,t)}{\partial t} = D_{eff} \cdot \nabla_r^2 f(r,t)$$

The time evolution of intensity profiles was simulated by solving the PDE system numerically using the R-package ReacTran (Soetaert and Meysman, 2012) that implements finite-difference grids. The radial axis from the spot center to the nucleus radius was split into 50 intervals to yield 50 concentric grid cells. A single ring-shaped grid cell was used for each radial interval assuming symmetry around the central spot position. Fluxes at the boundaries were set to zero. The model simulation resulted in radial profiles that were converted to averaged intensity values. The intensity in an area up to a radius of 3.5 µm for the pure diffusion model and from 0.0 to 1.0 µm for the reaction-diffusion model was averaged with the method described above for the image data.

**Parameter estimation from recovery curves**

We used individual recovery curves from off-spot FRAP measurements to estimate $D_{\mathrm{eff}}$ of the ligand constructs. The nuclear radius was determined from the segmented nuclear mask. The initial profile of free diffusible molecules $f(r,t=0)$ was estimated from the normalized profile of the first post-bleach frame fitted by a Gaussian with a plateau diameter of $r_{\mathrm{p}}$ and the parameter $\sigma$ describing the gaussian width. The amplitude was set to 1. Recovery of the normalized intensity in the bleach region was then fitted by a diffusion-only model with an immobile fraction using the *nls* function in R with multiple start values for the fit parameters. Starting values were varied between $D_{\mathrm{eff}}$ = 0.1 and 5 µm$^2$/s and an immobile fraction $f_{\mathrm{i}}$ = 0.1 and 0.5. The best fit out of all starting values was selected. The median of $D_{\mathrm{eff}}$ across single cell recovery curves for each ligand-target combination was used for further analysis. The normalized on-spot recovery curves were used to calculate $k_{\mathrm{off}}$ and immobile fraction of molecules at the binding site cluster. The immobile fraction $f_i$ was determined by fitting the data to a single exponential to the mostly binding dominated part of the recovery curve after 30 seconds:

$$I(t|t > 30s) = A + B \cdot \left(1 - e^{-k\cdot t}\right)$$

The immobile fraction was calculated as

$$f_i = 1 - A - B$$

with $f_{\mathrm{i}} \leq 0.5$. The full recovery time course was then fitted with the localized binding site cluster model with $D_{\mathrm{eff}}$ and $f_{\mathrm{i}}$ fixed. An approximated start value of the pseudo on-rate $\mathrm{k}_{\mathrm{on}}^{*}$ was calculated from the ratio of spot and nucleus intensity before bleaching.

$$spotRatio = \frac{median\left(I_{spot}(t < 0) - I_{spot\_bg}(t < 0)\right)}{median\left(I_{spot\_bg}(t < 0)\right)}$$

In (pre-bleach) equilibrium the ratio of bound and free molecules in the spot is given by

$c/f \ = \ k_{on}^{*}/k_{off}$, so that $\mathrm{k}_{\mathrm{on}}^{*}$ can be calculated as

$$k_{on}^{*} = \frac{c}{f} \cdot k_{off} = spotRatio \cdot k_{off}$$

A correction factor *bgRatio* that is multiplied with *spotRatio* was introduced as a free fit parameter to adjust for differences in the ratio of free and bound fraction. The initial profile $f(r,t=0)$ was estimated as described for the off-spot experiments and $c(r \leq r_s, t=0)$ was set to 0. Model simulations for a given parameter set yielded radial profiles of free and bound molecules $f(r,t)$ and $c(r,t)$ for each timepoint. These were processed and normalized like the imaging intensities:

$$y_{norm}(r,t) = (1 - f_i) \cdot \frac{f(r,t) + c(r,t)}{f(r,t = 240s) \cdot (1 + spotRatio \cdot bgRatio)}$$

This normalized profile was integrated from the spot center to 1.0 µm yielding a normalized time course $y_{norm}(t)$ that could be fitted to the data. The recovery curves were fitted by minimizing the sum of squared residual on a grid of parameter values for $k_{off}$ and *bgRatio* as described previously (Sprague et al., 2006). First, $k_{off}$ values were varied between $10^{-4}$ and 0.1 $s^{-1}$ in seven steps and *bgRatio* between 1 and 3.45 in steps of 0.35. The best parameter pair was used as a starting point for a refined optimization. In this second optimization the value of $k_{off}$ was multiplied by a factor between 0.2 and 8 and *bgRatio* was varied in steps of 0.03. The parameter pair with the smallest sum of squared residuals was selected as the best fit.

### Model for TF residence time dependent activation

TF residence time becomes functionally relevant if a kinetic proof-reading mechanism (Hopfield, 1974) is present that contains an energy-dissipating step subsequent to DNA binding like nucleosome remodeling (Shelansky and Boeger, 2020) or posttranslational modifications of the transcription complex or the TF itself (Kurosu and Peterlin, 2004). Such a mechanism in generic form is depicted in **Figure 6H** where the TF binds with rate constant $k_{on}$ to the free promoter (state *A*) and dissociates from the bound state *B* with rate constant $k_{off}$. An energy dependent step with rate $k_1$ leads to an activated TF bound state. The modified TF can dissociate from this state with the same dissociation rate constant $k_{off}$ as in state *B*. RNA is produced only from the activated state *C* with rate constant $k_t$ and is degraded with rate $k_m$. The total concentration of all promoter states is normalized to one, so that $A + B + C = 1$. The concentration of free TF is assumed to be high compared to the concentration of binding sites so that it can be absorbed into a pseudo-binding rate constant $k_{\text{on}}^* = k_{\text{on}} \cdot [TF]$. Furthermore, the loss of free modified TFs is taken to be comparatively fast so that rebinding of modified TFs can be neglected. The model is then described by the following system of ordinary differential equations:

$$\frac{dB}{dt} = k_{on}^* \cdot (1 - B - C) - (k_{off} + k_1) \cdot B$$

$$\frac{dC}{dt} = k_1 \cdot B - k_{off} \cdot C$$

$$\frac{dR}{dt} = k_t \cdot C - k_m \cdot R$$

The steady state levels are:

$$B = \frac{1}{\left(1 + \frac{k_{off}}{k_{on}^*}\right) \cdot \left(1 + \frac{k_1}{k_{off}}\right)}$$

$$C = \frac{1}{\left(1 + \frac{k_{off}}{k_{on}^*}\right) \cdot \left(1 + \frac{k_{off}}{k_1}\right)}$$

$$R = \frac{k_t}{k_m \cdot \left(1 + \frac{k_{off}}{k_{on}^*}\right) \cdot \left(1 + \frac{k_{off}}{k_1}\right)}$$

The TF concentration can be expressed in units of $K_D$ which leads to the equation for steady state RNA levels plotted in **Fig. 6 H, right**:

$$R = \frac{k_t}{k_m \cdot \left(1 + \frac{1}{[TF]}\right) \cdot \left(1 + \frac{k_{off}}{k_1}\right)}$$

The occupancy $\theta$ can be determined from the sum of (normalized) states *B* and *C*:

$$\theta = B + C = \frac{1}{1 + \frac{k_{off}}{k_{on}^*}} = \frac{[TF]}{[TF] + \frac{k_{off}}{k_{on}}}$$

Both steady state RNA levels and binding site occupancy depend on the TF concentration. The RNA levels are additionally limited by the last term of the denominator that contains the ratio of the TF modification rate constant and the dissociation rate. Hence, the residence time $\tau_{res} = 1/k_{off}$ regulates the steady state RNA level. This is illustrated by setting the modification rate constant to $k_1$ = 0.005 $s^{-1}$ and comparing two different dissociation rates $k_{off}$ = 0.006 $s^{-1}$ ($\tau_{res}$ = 167 s) and $k_{off}$ = 0.014 $s^{-1}$ ($\tau_{res}$ = 71 s). These $k_{off}$ or $\tau_{res}$ values reflect those observed for dCas9-GFP-VPR targeted to the *tet*O sites by mutated and wild type sgRNA. For simplicity the binding behavior was approximated by a weighted average of the apparent residence time and the immobile fraction $f_i$, that was assumed to have a residence time equal to the FRAP experiment duration ($\tau_{res}$ = 240 s).

$$k_{off} = \frac{1}{(1 - f_i) \cdot \tau_{FRAP} + f_i \cdot 240\ s}$$

The promoter becomes saturated at somewhat higher TF concentrations for the higher $k_{off}$ rate as computed for a value of $k_{on}$ = $10^5$ $M^{-1}$ $s^{-1}$, corresponding to $K_d$ = 60 nM and $K_d$ = 140 nM, respectively (**Figure 6H, left**). Notably, the RNA output is not only dependent on binding site occupancy but also directly reflects $k_{off}$. This is illustrated by the relation of RNA production and TF concentration given in units of the dissociation constant $K_d$ and thus normalized to the same promoter occupancy (**Figure 6H, right**). It can be seen that transcription increases with $\tau_{res}$ and the difference between the higher and lower $\tau_{res}$ persists even if full occupancy is reached. Thus, TF residence time and not binding site occupancy governs RNA production at saturating TF expression levels.

### Characterization of transcription incompetent BLInCR complexes

The BLInCR-dCas9 and BLInCR-loop complexes with VPR or VP16 were clearly enriched at the reporter array upon light induction (**Fig. S1A**). In addition, when coupled to VPR they efficiently induced BRD4 recruitment and H3K27 acetylation (**Fig. 5D, 5I**). Nevertheless, they were unable to induce transcription. While these complexes have a high turnover rate the same is true for the loop complexes for which AD binding does not involve a PHR-CIBN interaction (**Fig. 4, Table S5**). Thus, high TF turnover per se does not prevent efficient transcription activation. The BLInCR-dCas9 complex has been used previously to induce expression of selected single-copy genes, although with variable efficiency for the target genes and sgRNAs studied (Polstein and Gersbach, 2015). Furthermore, the BLInCR-TetR/rTetR complex is a strong transcription activator as shown here and in our previous work (Rademacher et al., 2017). To further dissect the inability of the BLInCR-dCas9 complex to induce transcription at the reporter array we conducted a number of additional experiments. We tested if the large size of the light-inducible dCas9 complexes (BLInCR-dCas9 VPR, 340 kDa; BLInCR-loop VPR, 348 kDa) hindered transcriptional induction. Accordingly, a fusion complex of comparable size (340 kDa) containing tetrameric GFP as a spacer between dCas9 and VPR (dCas9-GFP4-VPR) was studied (**Fig. S5A**). The GFP4 containing complex was a strong activator although induction of transcription was somewhat reduced as compared to dCas9-GFP-VPR (**Fig. S5A**). In contrast, a direct fusion of dCas9 with the catalytic core domain of the histone acetyl transferase p300 induced targeted H3K27 acetylation and BRD4 recruitment (**Fig. 5D**) but did not activate transcription (**Fig. S5A**). The dCas9-p300 fusion, however, is capable to activate certain single-copy genes (Hilton et al., 2015) including the *IL1RN* gene (Shrimp et al., 2018). *IL1RN* as well as *HBG1/2* were induced with CIBN-dCas9-CIBN (BLInCR-dCas9) in ref. (Polstein and Gersbach, 2015). Thus, it appears that promoters of some genes like *IL1RN* or *HBG1/2* are in a less repressed state than the reporter array used here, which has been shown to be H3K9-trimethylated and enriched in heterochromatin protein 1 (HP1) and the H3K9 methylase SUV39H1 (Janicki et al., 2004). Next, we tested if the BLInCR-dCas9 complex can activate a turboRFP reporter gene containing 5 x *tetO* sites upstream of a CMV minimal promoter in HeLa cells (**Fig. S5B**). This HeLa cell line was generated by random stable integration of a plasmid construct containing the silent 5xtetO-miniCMV-turboRFP reporter coupled to a cassette expressing a rTetR-transactivator (rTetR-TA) fusion that binds to the *tet*O sites in presence of doxycycline. Targeting a dCas9 VPR fusion to the *tet*O sites using a suitable sgRNA or recruitment of rTetR-TA by doxycycline addition efficiently induced the reporter expression after 24 hours (**Fig. S5C**). In contrast, targeting BLInCR-dCas9 with PHR-GFP-VP64 to the reporter as used in ref. (Polstein and

Gersbach, 2015) with a *tet*O sgRNA and blue-light illumination did not lead to turboRFP expression (**Fig. S5C, bottom**). Thus, we conclude that recruitment of the VPR activation domains consistently induces histone acetylation with all DBDs tested. Histone acetylation on its own is sufficient at some genes to subsequently induce transcription. However, the CMV minimal promoter used here appears to additionally require a certain configuration of the AD to initiate transcription that is not provided with BLInCR-dCas9/-loop constructs.

### Supplemental figures

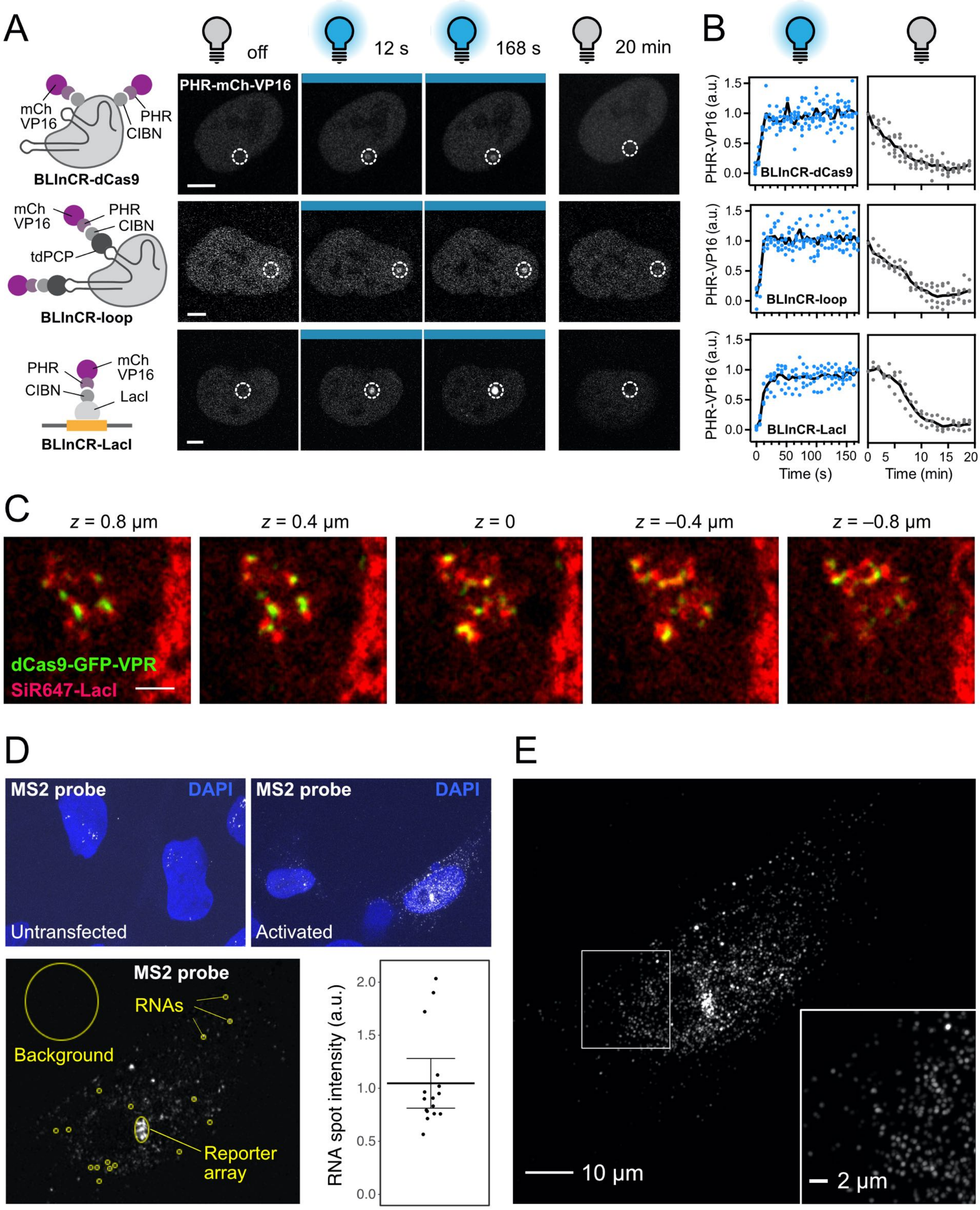


**Figure S1. Light induced activator binding and transcription activation**

(**A**) Confocal microscopy images of U2OS 2-6-3 cells expressing the BLInCR-dCas9, -loop and -LacI complexes depicted in the scheme. These constructs bind the reporter array upon blue light illumination and dissociate if light is switched off. The dashed circle marks the reporter array, which was identified by co-transfected TetR-YFP (not shown, see **Fig. 1 D**). Scale bars, 5 µm. (**B**) Quantification of the recruitment and dissociation kinetics in the presence or absence of blue light, respectively (*n* = 3 - 5 per construct). Solid line depicts the intensities averaged over all cells for each timepoint. (**C**) SRRF image *z*-stack of decondensed reporter array in a cell transfected with SiR647-labeled SNAPtag-LacI (red) and dCas9-GFP-VPR (green). Distance of *z*-slices, 0.4 µm; scale bar, 2 µm. (**D**) Single molecule RNA FISH of MS2-reporter RNA in U2OS 2-6-3 cells visualized by confocal microscopy. Top: Comparison of untransfected cells (left) and cells induced by transfection of CIBN-rTetR and PHR-GFP-VP16 (BLInCR rTetR) and overnight illumination. Bottom: Single *z*-plane image showing transcripts at the reporter array, in the nucleus and in the cytoplasm. About 80 nascent RNAs were detected as estimated by comparison with the intensity of single RNA spots (bottom, right) indicated with yellow circles (bottom, left). (**E**) SRRF maximum intensity projection image resolving a total of about ~2000 distinct RNA spots with ~1400 located in the nucleus and ~600 in the cytoplasm.

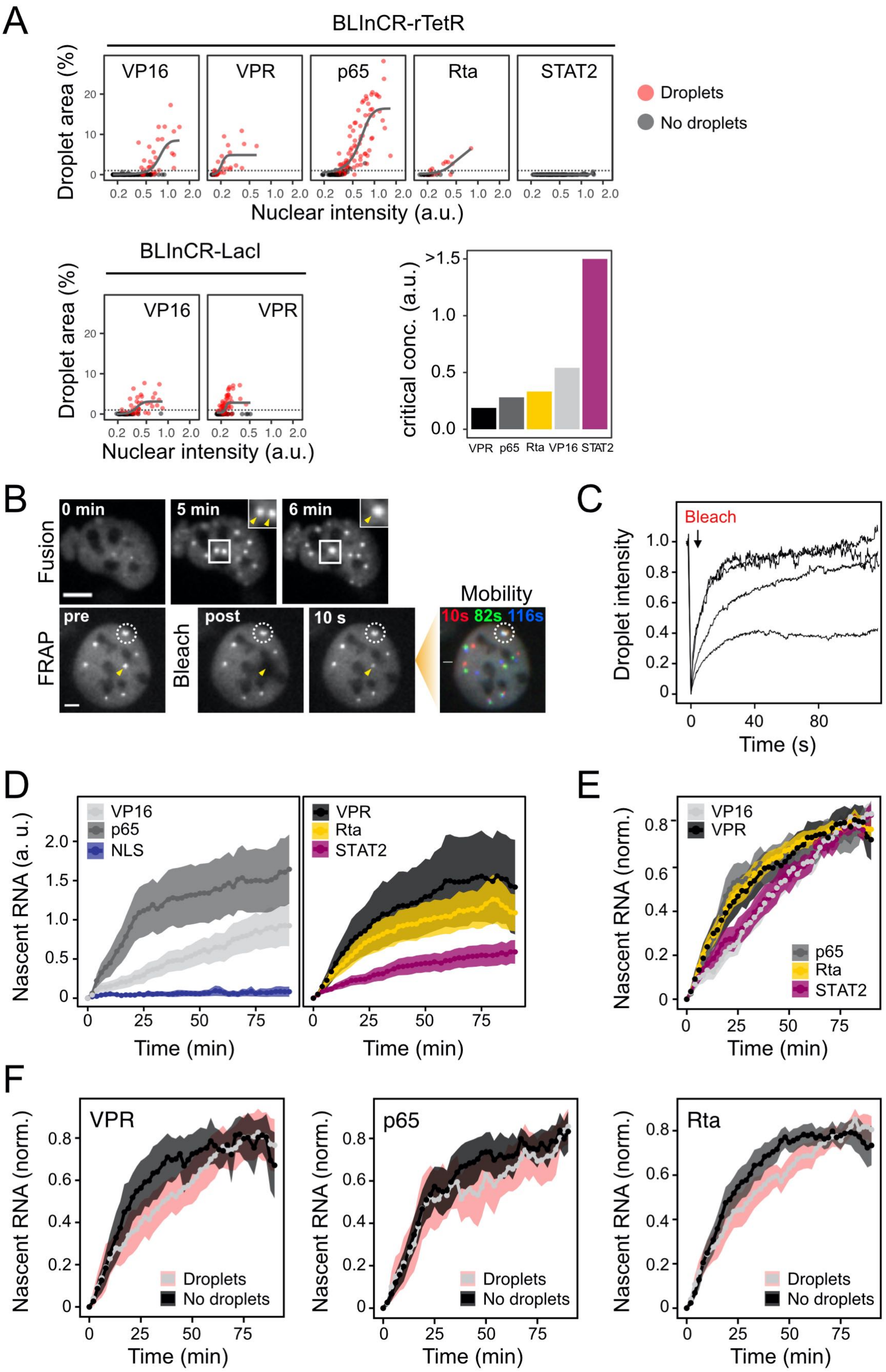


**Figure S2. AD droplet formation propensity and transcription activation kinetics**

(**A**) Optodroplet formation at different expression levels for the indicated PHR-GFP-AD construct in combination with different BLInCR targeting complexes. PHR-GFP-AD fluorescence was measured by microscopy in the nucleus after six cycles of illumination. Droplet abundance was quantitated as the area of segmented droplets in percent of the nuclear area as well as by manual annotation as droplet containing (red) or not (black) after visual inspection. The critical concentration was determined as the nuclear intensity at which the fitted logistic function (grey line) crossed the threshold at 1% (dashed line). (**B**) Liquid-like properties of VPR droplets formed outside the reporter array. Top: Image series showing the fusion of two droplets. Bottom: FRAP image series. Droplets were highly dynamic and recovered mostly within seconds after bleaching (yellow arrow). The droplets also showed displacement from their original position as apparent from their color-coded positions after 10, 82 and 116 s. The reporter array is marked by a dashed circle. Scale bars, 5 µm. (**C**) FRAP recovery curves of PHR-GFP-VPR optodroplets displaying predominantly fast recovery. (**D**) Averaged transcription activation kinetics of combined responder and non-responder cells (*n* = 37-132 cells per condition). Ribbon, 95% CI. (**E**) Averaged transcription activation kinetics after normalization to the maximum value of individual trajectories only for responding cells where nascent RNA was detectable. Ribbon, 95% CI. (**F**) Same as panel E for VPR, p65 and Rta but after dividing the cells into a group that displayed droplet formation outside the array and another that did not. Droplet formation did not enhance transcription activation kinetics. Ribbon, 95% CI; *n* = 13-40 cells per construct.

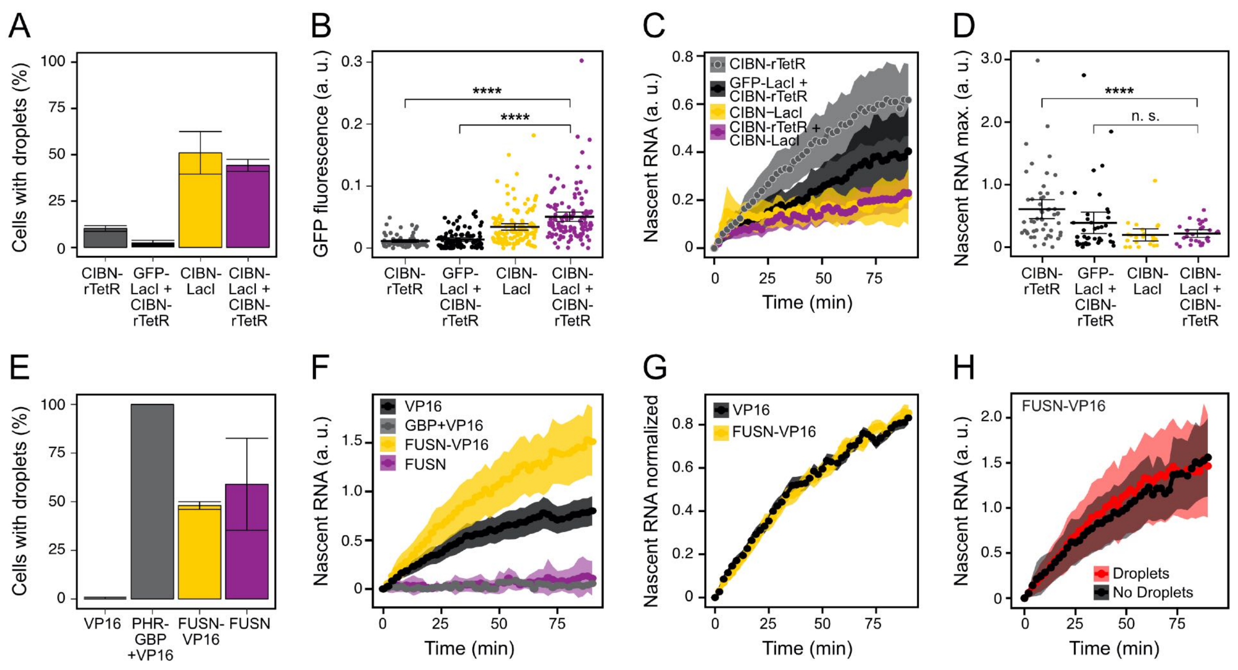


**Figure S3. Effect of droplet formation on transcription activation by VP16**

Light-induced transcription by VP16 recruited via BLInCR-rTetR was studied in dependence of factors that affected droplet formation of this activator construct. (**A**) Fraction of cells in induction experiments containing visible optodroplets; bar: mean, error bars: min. and max. of 2 replicate experiments. The presence of CIBN-LacI promoted droplet formation. (**B**) GFP signal at the reporter array. In the presence of CIBN-LacI additional PHR-GFP-VP16 molecules were recruited. Dots: single cell values, bar: mean, error bars: 95% CI. ****: $p < 0.0001$, two-sided Welch's t-test. (**C**) Average nascent RNA production time courses for responding cells. Addition of CIBN-LacI reduced transcription activation. Ribbon: 95% CI. **(D)** Nascent RNA plateau levels of responding cells. Dots: single cell values, bar: mean, error bars: 95% CI. n.s.: not significant, ****: $p < 0.0001$, two-sided Welch's t-test. **(E)** Fraction of cells displaying optodroplets. A fusion with FUSN or addition of PHR-GBP to bind an additional PHR domain enhanced droplet formation; bar: mean, error bars: min. and max. of 2 replicate experiments. **(F)** Average nascent RNA production time courses for responding cells. Ribbon: 95% CI. The FUSN-VP16 fusion enhanced transcription activation compared to VP16 only. (*n* = 94-134) **(G)** Average time courses of nascent RNA for responder cells normalized to maximum value of individual trajectories for comparison of activation kinetics of VP16 and FUSN-VP16. Transcription activation kinetics are not affected by the FUSN fusion. Ribbon, 95% CI. **(H)** Average time courses of nascent RNA production for responder cells activated by FUSN-VP16 divided into groups with or without visible optodroplets. No differences in activation kinetics and maximum levels between the two groups were apparent. Ribbon, 95% CI; *n* = 47 cells per condition.

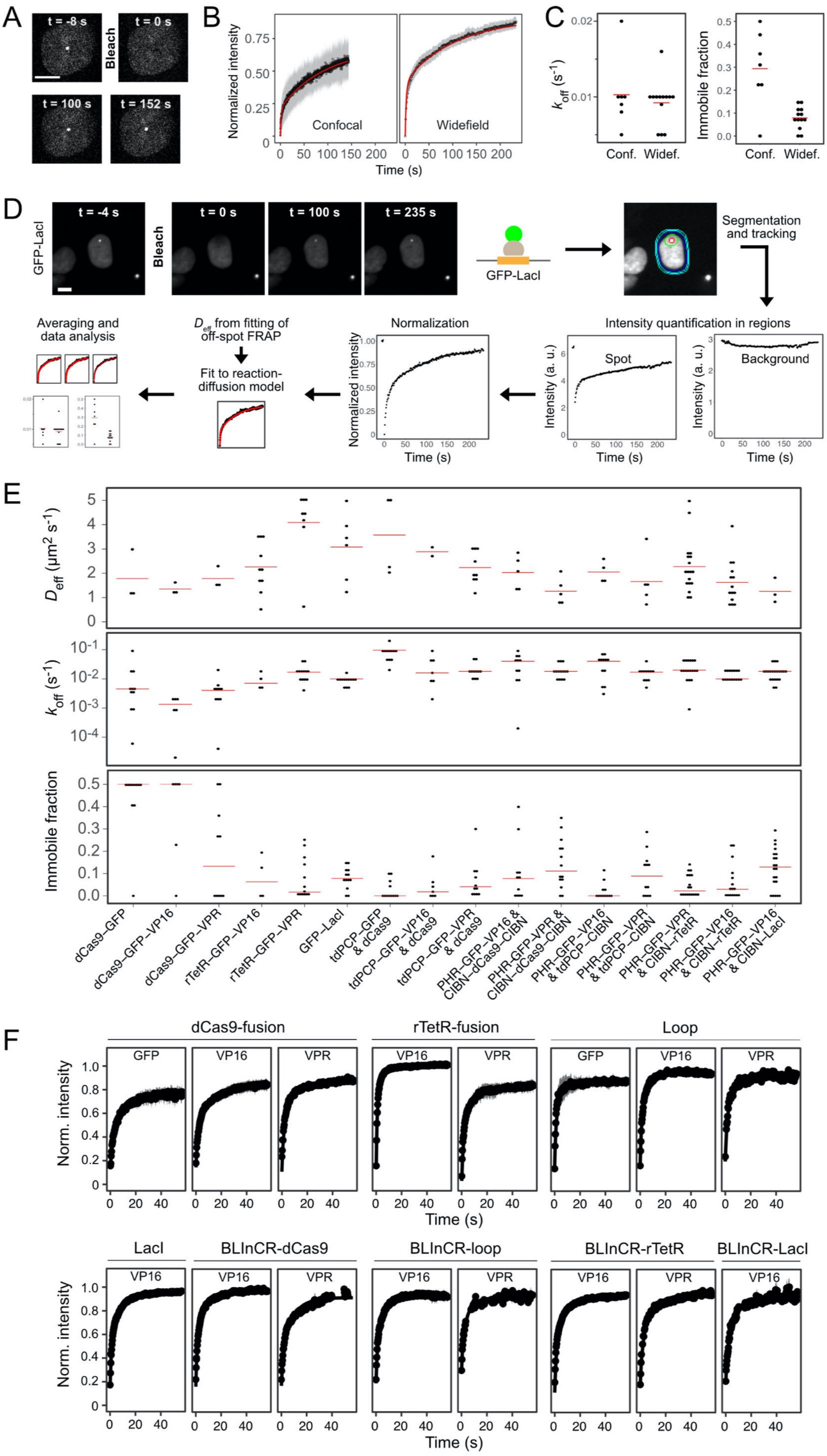
A
t = -8 s
t = 0 s
Bleach
t = 100 s
t = 152 s
B
Normalized intensity
Confocal
Widefield
Time (s)
C
k_off (s^-1)
Conf.
Widef.
Immobile fraction
D
GFP-LacI
t = -4 s
t = 0 s
t = 100 s
t = 235 s
Bleach
GFP-LacI
Segmentation and tracking
Intensity quantification in regions
Spot
Background
Intensity (a. u.)
Normalization
Normalized intensity
D_eff from fitting of off-spot FRAP
Fit to reaction-diffusion model
Averaging and data analysis
Time (s)
E
D_eff (μm² s^-1)
k_off (s^-1)
Immobile fraction
dCas9–GFP
dCas9–GFP–VP16
dCas9–GFP–VPR
rTetR–GFP–VP16
rTetR–GFP–VPR
GFP–LacI
tdPCP–GFP & dCas9
tdPCP–GFP–VP16 & dCas9
tdPCP–GFP–VPR & dCas9
PHR–GFP–VP16 & CIBN–dCas9–CIBN
PHR–GFP–VPR & CIBN–dCas9–CIBN
PHR–GFP–VP16 & tdPCP–CIBN
PHR–GFP–VPR & tdPCP–CIBN
PHR–GFP–VPR & CIBN–rTetR
PHR–GFP–VP16 & CIBN–rTetR
PHR–GFP–VP16 & CIBN–LacI
F
dCas9-fusion
rTetR-fusion
Loop
GFP
VP16
VPR
Norm. intensity
Time (s)
LacI
BLInCR-dCas9
BLInCR-loop
BLInCR-rTetR
BLInCR-LacI

**Figure S4. Experimental FRAP setup and data analysis**

(**A**) Image series of confocal FRAP of GFP-LacI bound to the reporter array. Scale bar: 10 µm. (**B**) Average recovery curves of GFP-LacI obtained by confocal and widefield FRAP. Ribbon, 95% CI; red line, fit of data to a reaction-diffusion model. The diffusive fraction is larger for widefield FRAP as discussed above in the Supplemental Methods section. (**C**) Binding parameters of GFP-LacI fits in confocal (Conf.) and widefield (Widef.) mode. Red bar: median. (**D**) Image analysis workflow for widefield FRAP illustrated for GFP-LacI as an example (scale bar 10 µm). Automated segmentation of spot (red), local background region (green), nucleus (blue) and background around nucleus (cyan) over the time course was followed by intensity quantification in these regions. The spot intensity was normalized, and binding parameters were obtained by fitting a reaction-diffusion model, which uses the effective diffusion coefficient determined in off-spot FRAP experiments. Normalized data, fit curves and fit parameters of single cells were averaged. (**E**) Distribution of parameters estimated from single cell recovery curves by a reaction-diffusion model. Effective diffusion coefficients were determined from off-spot FRAP while dissociation rate and immobile fraction were measured at the array. Red bar: median. (**F**) Average recovery curves for off-spot FRAP to determine the diffusion behavior of activation complexes. Fits of the data to a diffusion model are shown as solid line. Ribbon, 95% CI.

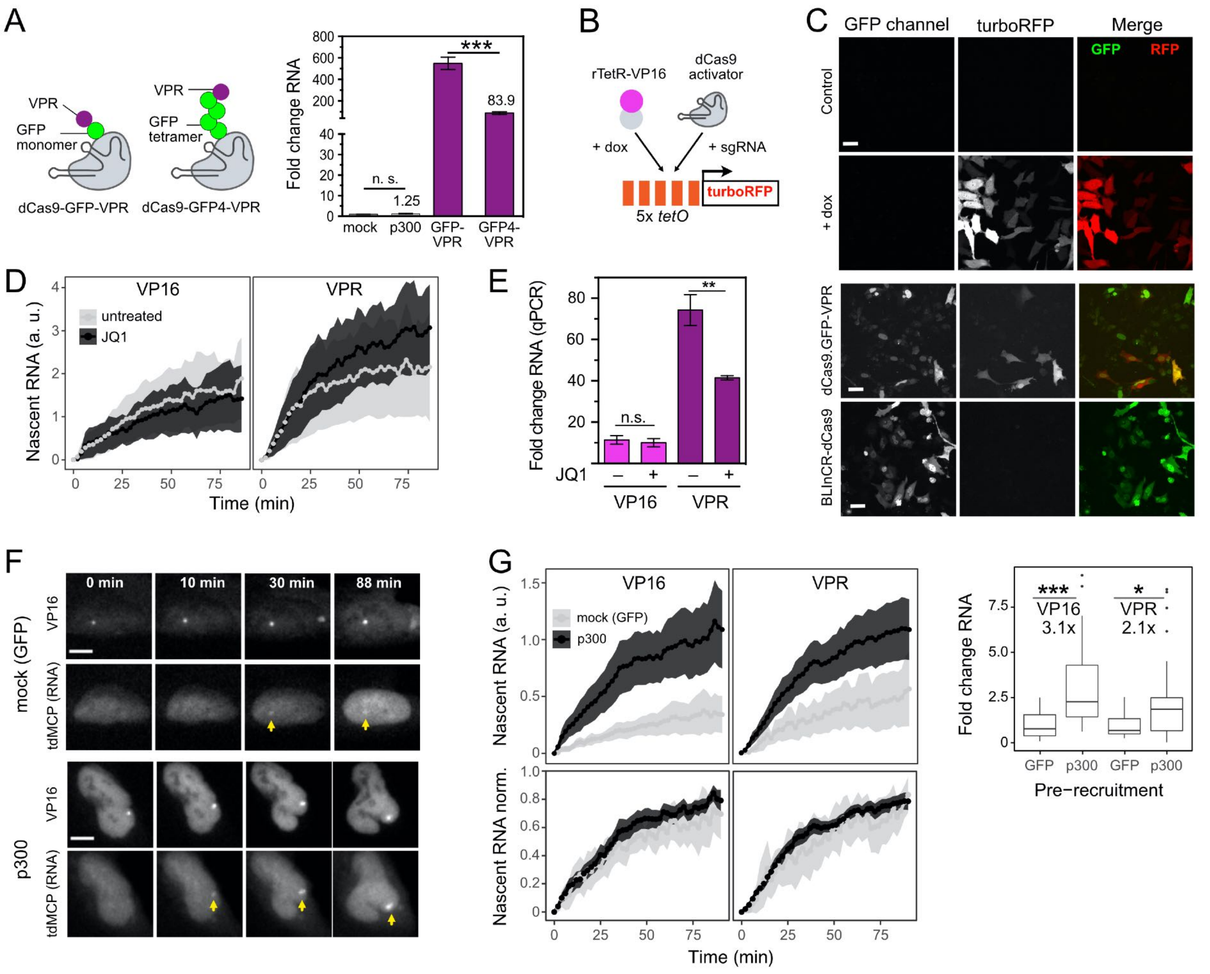

**Figure S5. Histone acetylation, BRD4 binding and transcription activation**

(**A**) Comparison of transcription induction measured by qRT-PCR of a dCas9-GFP4-VPR complex with a tetrameric GFP spacer and dCas9-GFP-VPR (from **Fig. 5A**) as a reference. Mean fold changes and standard deviation ($n$ = 3) of reporter RNA induction levels normalized to beta actin mRNA and relative to mock are shown; n. s., not significant, *, $p > 0.05$; ***, $p < 0.001$; two-sided unpaired Student's $t$-test. (**B**) Experimental strategy for testing the activation potential of the BLInCR-dCas9 complex in the HeLa 5x *tet*O-miniCMV-turboRFP reporter cell line stably expressing rTetR-VP16. The reporter turboRFP protein was induced by either addition of doxycycline to bind rTetR-VP16 at the *tet*O promoter sites or by targeting a dCas9 activator complex with sgRNA to this locus. (**C**) Fluorescence microscopy images of turboRFP reporter signal after 24 h of doxycycline induction of rTetR-VP16 or transient transfection with a *tet*O sgRNA and dCas9-GFP-VPR or the complex formed by CIBN-dCas9-CIBN and PHR-GFP-VP64 (BLInCR dCas9) as used previously (Polstein and Gersbach, 2015) and light illumination. While dCas9-GFP-VPR induced some transcription albeit at lower levels than rTetR-VP16, the BLInCR-dCas9 failed to do so. Scale bar: 20 μm. (**D**) Averaged time course data of light-induced nascent RNA production of VP16 and VPR BLInCR-rTetR for untreated cells or cells treated with JQ1 at 1 μM concentration starting 3 h prior to blue light illumination ($n$ = 7-20 cells per condition). Mean intensities with upper and lower boundaries corresponding to the 95% CI are shown. (**E**) Bulk reporter RNA levels measured by qRT-PCR for the conditions depicted in panel D at the 90 min endpoint of the time course. Data are fold changes in reporter RNA induction levels normalized to beta actin mRNA and relative to mock transfection ± s.d. ($n$ = 3); n.s., not significant, $p > 0.05$; **, $p < 0.01$ two-sided unpaired Student's $t$-test. (**F**) Representative widefield live cell images of time-resolved nascent RNA production (tdMCP-tdTomato) upon light-induced VP16 recruitment via BLInCR-rTetR. Cells either had dCas9-GFP ("mock") or dCas-GFP-p300core ("p300") pre-recruited to the *lac*O sites of the reporter before induction. Arrows indicate nascent RNA enriched at the reporter array. Scale bars, 10 µm. (**G**) Quantification of nascent RNA kinetics for the experimental setup described for panel (**F**) for VP16 and VPR ($n$ = 9-54 cells per condition). Pre-recruitment of p300 led to a 3.1-fold (VP16) or 2.1-fold (VPR) higher production of nascent RNA as shown in the box plot. Intensity values were normalized to the mean value of the respective mock pre-recruitment. *, $p < 0.05$; ***, $p < 0.001$; two-sided Welch's $t$-test. Bottom: Nascent RNA time courses normalized to their maximum show no effect of p300 pre-recruitment on the kinetics of the activation process.

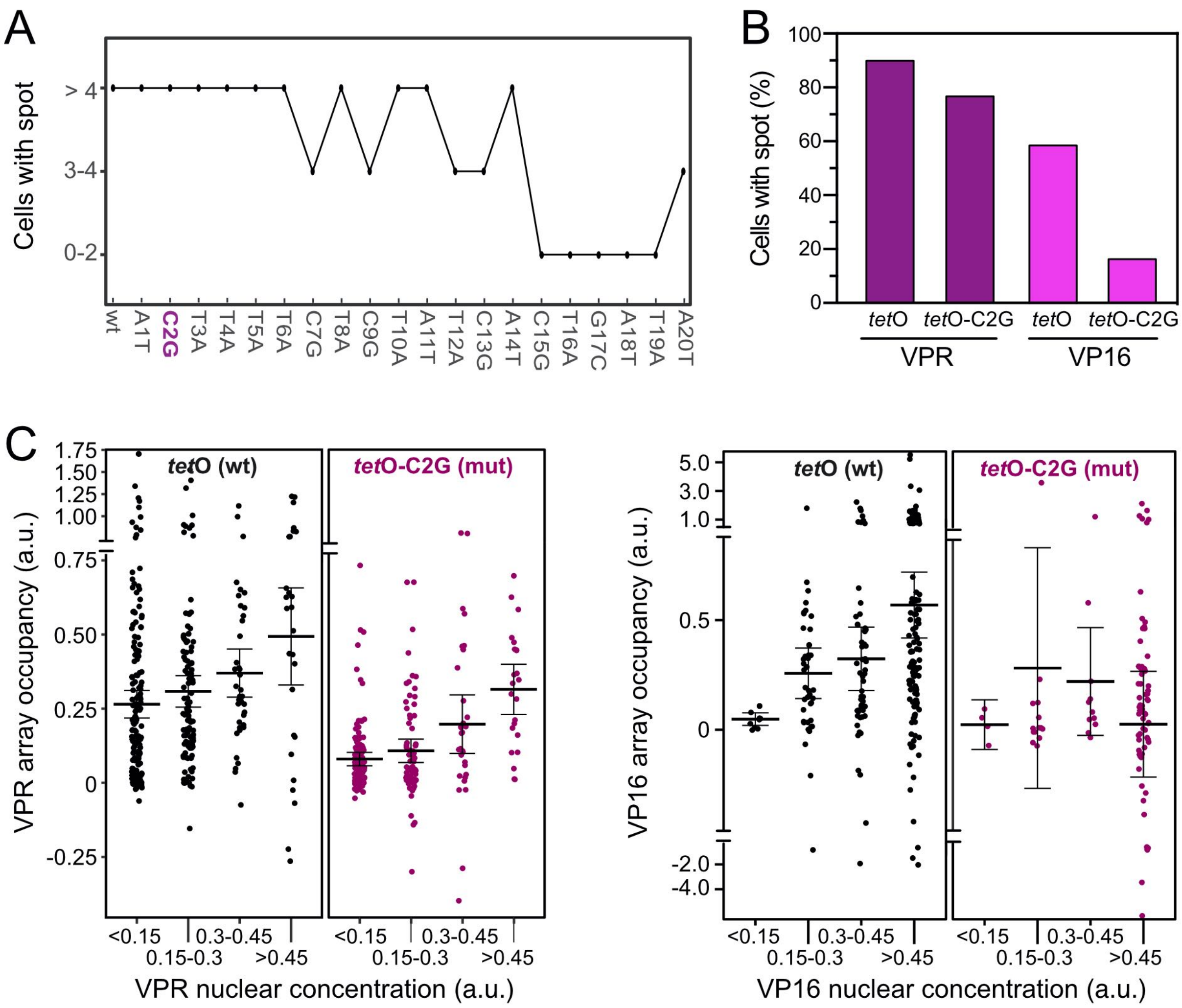


**Figure S6. Modulation of residence time via sgRNA mutations**

(**A**) Screen of *tet*O-sgRNA mutations that reduced dCas9 binding to *tet*O sites but were still enriched at the reporter array. The mutations introduced into the sgRNA targeting region are depicted on the x-axis. dCas9-GFP-VPR was recruited to the reporter with a given sgRNA and the number of cells with visible reporter spot recruitment (GFP) was counted per inspected region. (**B**) Fraction of cells with visible reporter array spots for dCas9-GFP-VPR or dCas9-GFP-VP16 recruited with the wildtype (*tet*O) or the mutated (*tet*O-C2G) sgRNA. A total of $n$ = 127-175 cells were evaluated per condition. (**C**) Reporter array occupancy in dependence of nuclear concentration of VPR or VP16 dCas9 fusion complexes with sgRNA-wt and sgRNA-mut. Concentrations were determined from nuclear GFP fluorescence intensities and cells were grouped according to this concentration. Occupancy corresponds to the GFP array intensity above background normalized to the co-transfected tagBFP-LacI array marker. Dots correspond to individual cells; mean and 95% CI error bars are indicated. Note the axis break to visualize the majority of cells and outliers in one plot.

### Supplemental Tables

#### Table S1. Plasmid constructs

| Plasmid | Comment | Reference |
|---|---|---|
| rTetR-GFP | Contains same NLS as VP16 constructs | This study |
| rTetR-GFP-VP16 | | This study |
| rTetR-GFP-VPR | | This study |
| CIBN-rTetR | Contains T2A-Puro resistance marker | This study |
| GFP-LacI | | (Jegou et al., 2009) |
| tagBFP-LacI | | Addgene #103839 (Rademacher et al., 2017) |
| CIBN-LacI | | Addgene #103814 (Rademacher et al., 2017) |
| SNAPtag-LacI | LacI in pSNAPf vector (New England Biolabs) | This study |
| dCas9 | dCas9 coding region from Addgene #60910, contains HA-tag | This study |
| dCas9-GFP | | (Erdel et al., 2020; Frank et al., 2021) |
| dCas9-GFP-VP16 | | This study |
| dCas9-GFP-VPR | | (Erdel et al., 2020; Frank et al., 2021) |
| dCas9-$GFP_4$-VPR | | This study |
| dCas9-GFP-p300 | p300 core domain from Addgene #61357 | This study |
| CIBN-dCas9-CIBN | | Addgene #60553 (Polstein and Gersbach, 2015) |
| PHR-GFP | Contains same NLS as VP16 constructs | (Rademacher et al., 2017) |
| PHR-GFP-VP16 | VP16 domain from (Gunther et al., 2013) | This study |
| PHR-GFP-VPR | VPR domain from Addgene #63798 | This study |
| PHR-GFP-p65 | p65 domain from Addgene #63798 | This study |
| PHR-GFP-Rta | Rta domain from Addgene #63798 | This study |
| PHR-GFP-STAT2 | STAT2 activation domain from (Frahm et al., 2006) | This study |
| PHR-GFP-FUSN | FUSN from Addgene #122148 (Bracha et al., 2018), | This study |
| PHR-GFP-FUSN-VP16 | FUSN from Addgene #122148 (Bracha et al., 2018), | This study |
| PHR-GBP | GBP from (Rothbauer et al., 2008) | This study |
| tdPCP-GFP | Tandem PCP from Addgene #40650 | This study |
| tdPCP-GFP-VP16 | TATA-box of the promoter removed | This study |
| tdPCP-GFP-VPR | TATA-box of the promoter removed | This study |
| tdPCP-CIBN | TATA-box of the promoter removed | This study |
| tdMCP-tdTomato | From Addgene #40649 and #54642 with TATA-box of the promoter removed | This study |
| mCherry-BRD4 | Murine BRD4 from ref. (Rafalska-Metcalf et al., 2010) | This study |

**Table S2. sgRNAs sequences used for dCas9 targeting**

| sgRNA | Targeting sequence (5'-3') |
|---|---|
| tetO-2xPP7 (wt) | GACTTTTCTCTATCACTGATA |
| tetO-2xPP7-A1T | GTCTTTTCTCTATCACTGATA |
| tetO-2xPP7-C2G | GAGTTTTCTCTATCACTGATA |
| tetO-2xPP7-T3A | GACATTTCTCTATCACTGATA |
| tetO-2xPP7-T4A | GACTATTCTCTATCACTGATA |
| tetO-2xPP7-T5A | GACTTATCTCTATCACTGATA |
| tetO-2xPP7-T6A | GACTTTACTCTATCACTGATA |
| tetO-2xPP7-C7G | GACTTTTGTCTATCACTGATA |
| tetO-2xPP7-T8A | GACTTTTCACTATCACTGATA |
| tetO-2xPP7-C9G | GACTTTTCTGTATCACTGATA |
| tetO-2xPP7-T10A | GACTTTTCTCAATCACTGATA |
| tetO-2xPP7-A11T | GACTTTTCTCTTTCACTGATA |
| tetO-2xPP7-T12A | GACTTTTCTCTAACACTGATA |
| tetO-2xPP7-C13G | GACTTTTCTCTATGACTGATA |
| tetO-2xPP7-A14T | GACTTTTCTCTATCTCTGATA |
| tetO-2xPP7-C15G | GACTTTTCTCTATCAGTGATA |
| tetO-2xPP7-T16A | GACTTTTCTCTATCACAGATA |
| tetO-2xPP7-G17C | GACTTTTCTCTATCACTCATA |
| tetO-2xPP7-A18T | GACTTTTCTCTATCACTGTTA |
| tetO-2xPP7-T19A | GACTTTTCTCTATCACTGAAA |
| tetO-2xPP7-A20T | GACTTTTCTCTATCACTGATT |
| lacO-2xPP7 (wt) | GTCCGCTCACAATTCCACATG |
| tetO-turboRFPreporter-2xPP7 | GATACGTTCTCTATCACTGAT |

All sgRNAs were cloned into the U6 promoter-driven sgRNA expression vector originally derived from Addgene #61424 and engineered to contain two PP7 stem loops PP7. The PP7 loop sequence was adapted from ref. (Zalatan et al., 2015).

**Table S3. Propensity of the activation domain to form optodroplets**

| PHR-GFP-AD | DNA binder | Cell number | Droplets (%)[a] | $I_{crit}$ (a. u.) [b] |
|---|---|---|---|---|
| VP16 | CIBN-rTetR | 131 | 29 | 0.54 |
| VPR | CIBN-rTetR | 38 | 86 | 0.19 |
| p65 | CIBN-rTetR | 129 | 72 | 0.28 |
| Rta | CIBN-rTetR | 34 | 41 | 0.33 |
| STAT2 | CIBN-rTetR | 103 | 0 | >1.5 [c] |
| VP16 | CIBN-LacI | 79 | 56 | 0.34 |
| VPR | CIBN-LacI | 106 | 63 | 0.23 |

Cells were classified as positive for droplet formation if they displayed nuclear optodroplets in microscopy images in addition to the signal at the reporter array (**Fig. 1E**).

[a] The percentage of cells with droplets depends on the nuclear concentration range for each construct, but allows a simple distinction between droplet-forming and non-droplet-forming ADs at typical expression levels.

[b] The critical value for droplet formation $I_{crit}$ was determined from the relation of nuclear PHR-GFP-AD concentration and droplet abundance shown in **Fig. S2A**.

[c] If droplet abundance did not exceed the threshold value within the measured nuclear concentrations the critical concentration is reported as greater than the highest observed nuclear concentration.

**Table S4. Transcription activation kinetics**

| PHR-GFP-AD | Condition | Cell number | Respon-ders (%) | $t_{1/2}$ (min) [a] | Maximum RNA value (a. u.) [a] |
|---|---|---|---|---|---|
| VP16 | All cells | 64 | 67 | 42 (37-46) | 1.2 (0.89-1.6) |
| VPR | All cells | 37 | 84 | 28 (23-33) | 1.7 (1.1-2.4) |
| VPR | Cells without droplets | 15 | 87 | 25 (17-34) | 1.1 (0.83-1.4) |
| VPR | Cell with droplets | 22 | 82 | 30 (24-36) | 2.2 (1.0-3.3) |
| p65 | All cells | 52 | 67 | 26 (21-31) | 2.1 (1.6-2.6) |
| p65 | Cells without droplets | 23 | 78 | 26 (19-33) | 2.3 (1.5-3.0) |
| p65 | Cell with droplets | 29 | 59 | 26 (18-34) | 1.9 (1.2-2.7) |
| Rta | All cells | 77 | 92 | 28 (25-31) | 1.2 (0.92-1.5) |
| Rta | Cells without droplets | 33 | 94 | 25 (21-29) | 1.2 (0.73-1.7) |
| Rta | Cell with droplets | 44 | 91 | 31 (26-36) | 1.2 (0.85-1.6) |
| STAT2 | All cells | 132 | 42 | 38 (34-43) | 0.95 (0.66-1.2) |
| **Droplet induction experiments** | | | | | |
| VP16 | No additional factors | 74 | 70 | 35 (30-39) | 0.61 (0.46-0.76) |
| VP16 | GFP-LacI | 97 | 42 | 34 (28-39) | 0.39 (0.22-0.56) |
| VP16 | CIBN-LacI | 118 | 24 | 31 (23-38) | 0.22 (0.16-0.28) |
| VP16 | CIBN-LacI but no CIBN-rTetR | 126 | 18 | 20 (13-28) | 0.20 (0.09-0.29) |
| VP16 | No additional factors | 154 | 84 | 34 (31-36) | 0.78 (0.63-0.93) |
| VP16 | PHR-GBP | 24 | 4 | n. d. [b] | n. d. [b] |
| FUS-VP16 | No additional factors | 108 | 87 | 37 (34-41) | 1.5 (1.1-1.8) |
| FUS | No additional factors | 57 | 5 | n. d. [b] | n. d. [b] |

The RNA production at the reporter gene cluster was followed over time via the tdMCP-tdTomato signal. CIBN-rTetR was used as DBD module unless stated otherwise.

[a] The maximum of RNA produced was determined from the last five time points at the plateau of the single cell time course, and the time $t_{1/2}$ was determined where half of this values was reached. Mean values and 95% CIs were calculated from the analysis of responding cells that showed an RNA signal at the reporter array. Data for VP16 and p65 as well as for VPR, Rta and STAT2 were acquired together. A direct comparison of VPR and VP16 done in other experiments yielded a VPR/VP16 ratio of maximum activation values of ~1.5 after 90 minutes.

[b] Values could not be determined due to the low number of responder cells.

**Table S5. FRAP parameters of TF dynamics**

| Protein(s) | DNA | Residence time (s) | Immobile fraction (%) | $D_{eff}$ (μm$^2$/s) | *n* |
|---|---|---|---|---|---|
| GFP-LacI | *lac*O | 108 (91-134) | 8 (5-11) | 2.3 (1.5-3.0) | 13 |
| GFP-LacI (confocal) | *lac*O | 97 (69-167) | 29 (14-45) [b] | 3.3 (2.0-4.5) | 7 |
| dCas9-GFP | *lac*O | 74 (32->240) | 44 (34-54) | 1.8 (0-4.3) | 11 |
| dCas9-GFP-VP16 | *lac*O | >240 (>240) | 37 (15-59) | 1.4 (0.8-2.0) | 6 |
| dCas9-GFP-VPR | *lac*O | 204 (112->240) | 19 (4-34) | 1.8 (0.7-2.9) | 10 |
| dCas9-GFP-VPR | *tet*O | 124 (75->240) | 36 (25-47) | 0.6 (0.4-0.9) | 10 |
| dCas9-GFP-VPR | *tet*O-C2G | 57 (34-184) | 7 (0-16) | - [a] | 7 |
| rTetR-GFP-VP16 | *tet*O | 132 (74->240) | 5 (0-14) | 4.3 (3.0-5.4) | 6 |
| rTetR-GFP-VPR | *tet*O | 57 (42-86) | 7 (2-12) | 3.1 (1.6-4.5) | 13 |
| dCas9 + tdPCP-GFP | *lac*O | 12 (9-18) | 3 (0-5) | 3.4 (0.9-6.2) | 12 |
| dCas9 + tdPCP-GFP-VP16 | *lac*O | 33 (17->240) | 4 (0-10) | 2.9 (0.6-5.2) | 7 |
| dCas9 + tdPCP-GFP-VPR | *lac*O | 47 (33-83) | 7 (1-13) | 2.2 (1.7-2.8) | 11 |
| CIBN-dCas9-CIBN + PHR-GFP-VP16 | *lac*O | 28 (18-58) | 10 (1-19) | 2.0 (1.2-2.9) | 11 |
| CIBN-dCas9-CIBN + PHR-GFP-VPR | *lac*O | 49 (37-72) | 14 (8-21) | 1.3 (0.6-1.9) | 14 |
| tdPCP-CIBN + dCas9 PHR-GFP-VP16 | *lac*O | 29 (20-53) | 2 (0-4) | 2.1 (1.4-2.8) | 12 |
| tdPCP-CIBN + dCas9 PHR-GFP-VPR | *lac*O | 60 (45-91) | 10 (3-16) | 1.7 (0.4-3.0) | 12 |
| CIBN-rTetR + PHR-GFP-VP16 | *tet*O | 42 (33-58) | 4 (2-7) | 2.3 (1.7-2.8) | 16 |
| CIBN-rTetR + PHR-GFP-VPR | *tet*O | 71 (60-88) | 6 (2-10) | 1.6 (1.1-2.1) | 16 |

Measurements were conducted with the FRAP widefield microscopy setup except for the indicated measurement of GFP-LacI on a confocal microscopy. Mean values and 95% confidence intervals in brackets were determined as described in the Supplemental Methods section.

[a] Not determined. For fitting of the diffusion-binding model the effective diffusion coefficient for dCas9-GFP-VPR with *tet*O-sgRNA(wt) was used.

[b] This value is expected to differ from the corresponding value in the widefield setup since less freely diffusing fluorescent particles above and below the array are visible in a confocal setup.

**Table S6. Reporter RNA expression measured by qRT-PCR**

| DNA binder | AD | Treatment | RNA fold-change |
|---|---|---|---|
| – (mock transfection, reference for normalization) | – | 24 h light | 1.0 |
| – (untransfected) | – | 24 h light | 0.9 |
| dCas9-GFP-VP16 | VP16 (fusion) | 24 h light | 6.4 |
| dCas9-GFP-VPR | VPR (fusion) | 24 h light | 550 |
| dCas9-$GFP_4$-VPR | VPR (fusion) | 24 h light | 84 |
| dCas9-GFP-VP16 (tetO-C2G) | VP16 (fusion) | 24 h light | 0.9 |
| dCas9-GFP-VPR (tetO-C2G) | VPR (fusion) | 24 h light | 31 |
| dCas9-GFP-p300 | p300 (fusion) | 24 h light | 1.3 |
| rTetR-GFP-VP16 | VP16 (fusion) | 24 h light + dox | 35 |
| rTetR-GFP-VPR | VPR (fusion) | 24 h light + dox | 217 |
| dCas9 (tetO-PP7) | tdPCP-GFP-VP16 | 24 h light | 4.0 |
| dCas9 (tetO-PP7) | tdPCP-GFP-VPR | 24 h light | 490 |
| CIBN-dCas9-CIBN | PHR-GFP-VP16 | 24 h light | 1.7 |
| CIBN-dCas9-CIBN | PHR-GFP-VPR | 24 h light | 1.8 |
| CIBN-dCas9-CIBN | PHR-GFP-VP16 | dark | 1.3 |
| CIBN-dCas9-CIBN | PHR-GFP-VPR | dark | 1.6 |
| dCas9 + tdPCP-CIBN | PHR-GFP-VP16 | 24 h light | 1.0 |
| dCas9 + tdPCP-CIBN | PHR-GFP-VPR | 24 h light | 1.4 |
| CIBN-rTetR | PHR-GFP-VP16 | dox, 24 h light | 32 |
| CIBN-rTetR | PHR-GFP-VPR | dox, 24 h light | 17.5 |
| CIBN-rTetR + CIBN-LacI | PHR-GFP-VP16 | dox, 90 min light | 3.1 |
| CIBN-rTetR + GFP-LacI | PHR-GFP-VP16 | dox, 90 min light | 5.3 |
| CIBN-LacI | PHR-GFP-VP16 | dox, 90 min light | 1.7 |
| CIBN-rTetR | PHR-GFP-VP16 | dox, 90 min light | 7.0 |
| CIBN-rTetR | PHR-GFP-FUSN | dox, 90 min light | 1.8 |
| CIBN-rTetR | PHR-GFP-FUSN-VP16 | dox, 90 min light | 70 |
| CIBN-rTetR | PHR-GFP-VP16 + PHR-GBP | dox, 90 min light | 1.9 |
| CIBN-rTetR [a] | PHR-GFP-VP16 | dox, untreated, 90 min light | 11 |
| CIBN-rTetR [a] | PHR-GFP-VPR | dox, untreated, 90 min light | 74 |
| CIBN-rTetR [b] | PHR-GFP-VP16 | dox, 3 h JQ1, 90 min light | 10 |
| CIBN-rTetR [b] | PHR-GFP-VPR | dox, 3 h JQ1, 90 min light | 41 |
| CIBN-rTetR [c] | PHR-GFP-VP16 | dCas9-GFP *lac*O, dox, 90 min light | 9.7 |
| CIBN-rTetR [c] | PHR-GFP-VPR | dCas9-GFP *lac*O, dox, 90 min light | 111 |
| CIBN-rTetR [c] | PHR-GFP-VP16 | dCas9-GFP-p300 *lac*O + dox, 90 min light | 15 |
| CIBN-rTetR [c] | PHR-GFP-VPR | dCas9-GFP-p300 *lac*O + dox, 90 min light | 95 |

Doxycycline (dox) was added directly after transfection and cells were illuminated for 24h for the “24h light” experiments. For the “90 min light” experiments dox was added 24h after transfection; cells were exposed to light after 15 min for 90 min. JQ1 treatment started 3 h before the start of illumination. RNA levels were normalized to beta actin mRNA for each sample and fold-changes were determined from the average of three measurements relative to the mock transfected cells.

[a] Reference for the JQ1 treatment experiment.

[b] Cells were treated with JQ1 at a 1 μM concentration in the dark for 3 h and then activated with light.

[c] Histone hyperacetylation was induced by recruiting dCas9-GFP-p300 with the *lac*O sgRNA for 24 h.

**Table S7. Histone acetylation, BRD4 binding and transcription activation**

| DNA binder and readout | PHR-GFP-AD | Condition | Cell number | Respon-ders (%) | $t_{1/2}$ (min) | Maximum value (a. u.) [a] |
|---|---|---|---|---|---|---|
| dCas9 + tdPCP-CIBN, mCherry-BRD4 | VP16 | – | 37 | 27 | 13 (7-20) | 0.007 (0.005-0.010) |
| | | JQ1 | 85 | 0 | – | 0.002 (0.002-0.003) |
| | VPR | – | 13 | 92 | 13 (8-18) | 0.024 (0.016-0.032) |
| | | JQ1 | 10 | 10 | – | 0.005 (0.002-0.008) |
| CIBN-rTetR, tdMCP-tdTomato (RNA) | VP16 | – | 29 | 69 | 35 (30-39) | 1.7 (0.91-2.5) |
| | | JQ1 | 12 | 58 | 28 (18-39) | 1.4 (0.63-2.2) |
| | VPR | – | 15 | 87 | 29 (20-39) | 2.1 (1.0-3.2) |
| | | JQ1 | 21 | 81 | 33 (27-38) | 3.0 (2.0-4.0) |
| | VP16 | dCas9-GFP | 52 | 25 | 35 (25-44) | 0.35 (0.18-0.51) |
| | | dCas9-GFP-p300 | 49 | 53 | 31 (27-36) | 1.1 (0.76-1.4) |
| | VPR | dCas9-GFP | 27 | 33 | 37 (19-54) | 0.51 (0.23-0.79) |
| | | dCas9-GFP-p300 | 72 | 75 | 31 (27-34) | 1.1 (0.82-1.4) |

Transcription activation time course parameters were determined as described for **Table S4**.

[a] Maximum values for RNA production can be compared directly only for the same DNA binding module and experimental conditions. For the experiments with BRD4, values are given for the whole population of responding and non-responding cells due to the low number of responding cells detected after JQ1 treatment.

**Table S8. Binding site occupancy of dCas9**

| DNA binder | AD | sgRNA | Occupancy [a] | Number of cells | Visible array (%) [b] |
|---|---|---|---|---|---|
| dCas9 | tdPCP-GFP-VP16 | *tet*O-2xPP7 (wt) | 0.53 (0.41-0.66) | 166 | 78 |
| dCas9 | tdPCP-GFP-VPR | *tet*O-2xPP7 (wt) | 0.99 (0.77-1.22) | 164 | 93 |
| dCas9-GFP-VP16 | VP16 (fusion) | *tet*O-2xPP7 (wt) | 0.24 (0.17-0.30) | 138 | 59 |
| | | *tet*O-2xPP7-C2G (mut) | 0.03 (0.01-0.06) | 127 | 17 |
| dCas9-GFP-VPR | VPR (fusion) | *tetO*-2xPP7 (wt) | 0.32 (0.26-0.37) | 175 | 90 |
| | | *tet*O-2xPP7-C2G (mut) | 0.12 (0.09-0.14) | 163 | 76 |

Binding site occupancy at the reporter array were determined as the ratio of the GFP fluorescence of the activator complex and the blue fluorescence signal of tagBFP-LacI as an array marker.

[a] Mean value and 95% CI. Data can be directly compared only for experiments conducted with the same DNA binding module.

[b] Fraction of cells that had the activator complex GFP signal enriched at the site of the array marked by tagBFP-LacI.
